# Cotranslational folding cooperativity of contiguous domains of α-spectrin

**DOI:** 10.1101/653360

**Authors:** Grant Kemp, Ola B. Nilsson, Pengfei Tian, Robert B. Best, Gunnar von Heijne

## Abstract

Proteins synthesized in the cell can begin to fold during translation before the entire polypeptide has been produced, which may be particularly relevant to the folding of multidomain proteins. Here, we study the cotranslational folding of adjacent domains from the cytoskeletal protein α-spectrin using Force Profile Analysis (FPA). Specifically, we investigate how the cotranslational folding behavior of the R15 and R16 domains are affected by their neighboring R14 and R16, and R15 and R17 domains, respectively. Our results show that the domains impact each other’s folding in distinct ways that may be important for the efficient assembly of α-spectrin, and may reduce its dependence on chaperones. Furthermore, we directly relate the experimentally observed yield of full-length protein in the FPA assay to the force exerted by the folding protein in pN. By combining pulse-chase experiments to measure the rate at which the arrested protein is converted into full-length protein with a Bell model of force-induced rupture, we estimate that the R16 domain exerts a maximal force on the nascent chain of ∼15 pN during cotranslational folding.

**Significance:** In living cells, proteins are produced in a sequential way by ribosomes. This vectoral process allows the growing protein chain to start to fold before translation has been completed. Thereby, cotranslational protein folding can be significantly different than the folding of a full-length protein in isolation. Here we show how structurally similar repeat domains, normally produced as parts of a single long polypeptide, affect the cotranslational folding of their neighbors. This provides insight into how the cell may efficiently produce multidomain proteins, paving the way for future studies *in vivo* or with chaperones. We also provide an estimated magnitude of the mechanical force on the nascent chain generated by cotranslational folding, calculated from biochemical measurements and molecular dynamics simulations.

## Introduction

Classically, protein folding has been studied *in vitro* using purified proteins, providing a detailed picture of how the biophysical properties of a given polypeptide chain affect folding pathways. However, studies of purified proteins do not reflect the vectorial nature of biological protein synthesis, where each amino acid is added sequentially on a time scale that allows the growing protein to explore its continually expanding energy landscape (1).

The discovery (2, 3) and engineering (4, 5) of translational arrest peptides (APs) has provided us with a new tool to examine how proteins fold cotranslationally. To date, the folding of several small proteins and protein domains has been studied using the force-sensitive AP from the *E. coli* SecM protein, establishing a method we have called Force-Profile Analysis (FPA) (6–15). Recently, we demonstrated that FPA faithfully picks up cotranslational protein folding events observed by direct biophysical measurements (15).

Spectrin, the primary structural component of the cellular cytoskeleton (16–19), is composed of two heterodimers of α- and β-spectrin that come together in a coiled tetramer (20). The 285 kDa non-erythroid αII isoform of spectrin contains 20 three-helix bundle domains, which provide interesting model systems for protein folding. This is because, while they are structurally nearly identical (21), they vary in stability (22–25), folding rate (26, 27), and folding mechanism (28). In particular, folding of the 15^th^, 16^th^, and 17^th^ α-spectrin domains (herein called R15, R16, and R17, respectively) has been extensively studied *in vitro* (24, 27, 29–32).

Previously, using FPA, we demonstrated that R15 and R16 are able to fold cotranslationally and that the point at which this folding begins does not seem to be influenced by the *in vitro* folding rate (9). Specifically, while R15 folds very quickly *in vitro*, cotranslational folding of R16 starts at shorter chain lengths than does folding of R15. Indeed, even when the folding core of R15 was substituted into R16, which gives R16 the folding properties of R15 *in vitro* (33), the protein was still able to fold earlier than R15 during translation. When the minimal, five-residue R15 folding nucleus (34) was substituted into R16, this led to late cotranslational folding, but early folding was restored when this mutant was expressed in tandem with R15, as it would be *in vivo*. This demonstrated that the mutant R16 variants fold cotranslationally via early intrachain contacts, at chain lengths where the R15 folding nucleus cannot yet form. This was not observed *in vitro* because the purified mutant R16 protein would preferentially fold using the “fast” R15 nucleus rather than the slower R16 contacts. Cotranslationally, R16 folds using the first available pathway, i.e., the slower R16 folding contacts.

Here, we have extended our studies on spectrin to determine how the presence of neighboring domains affect the folding of R15 and R16. We find that the cotranslational folding of these two domains is differently affected by their respective up- and downstream neighbor domains, a previously unappreciated feature of spectrin folding. We were also able to observe that the synthesis of native, likely helical, structures downstream of the folding spectrin domain increases the observed folding force and further stabilizes the fold even in the presence of amino acid substitutions that destabilize the final native spectrin structure.

While it has been shown previously that the cotranslational folding of proteins can provide a mechanical force to release SecM-mediated ribosomal stalling (6), the magnitude of the force has never been directly measured by FPA. Using pulse-chase experiments, we have now measured the rate at which arrested protein is converted to full-length protein under standard experimental conditions. These rates were fit to a Bell model (35) for force-induced rupture, using parameters elucidated by molecular dynamics simulations (12), to provide an estimate of the magnitude of the pulling force exerted on the nascent chain by cotranslational protein folding. The determined force is in good agreement with previously published measurements using optical tweezers (6).

## Results

### The force profile analysis (FPA) assay

APs are short stretches of polypeptide that interact with the ribosome exit tunnel in such a way that translation is stalled when the last codon in the portion of the mRNA that codes for the AP is located in the ribosomal A-site (36). APs generally have regulatory functions in cells, controlling, *e.g*., translation initiation on downstream ORFs in polycistronic mRNAs (36). Interestingly, the stalling efficiency of many APs has been shown to be sensitive to pulling forces exerted on the nascent polypeptide chain, with high pulling force leading to reduced stalling (6, 37, 38). APs can therefore be used as force sensors, reporting on cotranslational events that in one way or another generate pulling forces on the nascent chain (7, 38, 39). The factors influencing such force-generating events were recently examined using molecular simulations and statistical mechanics models, in particular the effect of translation speed (40). Although arrest peptide measurements of force do not include the effect of translation speed directly, this could be accounted for by using a suitable kinetic model for coupled folding and translation (41).

In FPA, a force-generating domain in a polypeptide is placed at increasing distances upstream of an AP, Fig. 1a, and the degree of translational stalling is measured for the corresponding series of protein constructs. Figure 1b shows how FPA can be applied to study the cotranslational folding of soluble protein domains, *e.g*., tandem spectrin domains. In the construct shown on the left, the chain is long enough for one of the two spectrin domains to have already folded, while the second spectrin domain is largely buried in the exit tunnel and thus cannot fold when the ribosome reaches the C-terminal end of the AP. Therefore, little force is generated on the AP, and stalling is efficient. In the right-hand construct, the chain is so long that both spectrin domains have folded prior to the ribosome reaching the C-terminal end of the AP, and again little force is generated on the AP. In the middle construct, however, the chain is just long enough that the second spectrin domain can begin to fold if the linker that connects it to the AP is stretched beyond its equilibrium length. Under this condition, some of the free energy gained upon folding will be converted to tension in the linker and generate a pulling force on the AP, resulting in reduced stalling and increased production of full-length protein.

**Figure 1.**
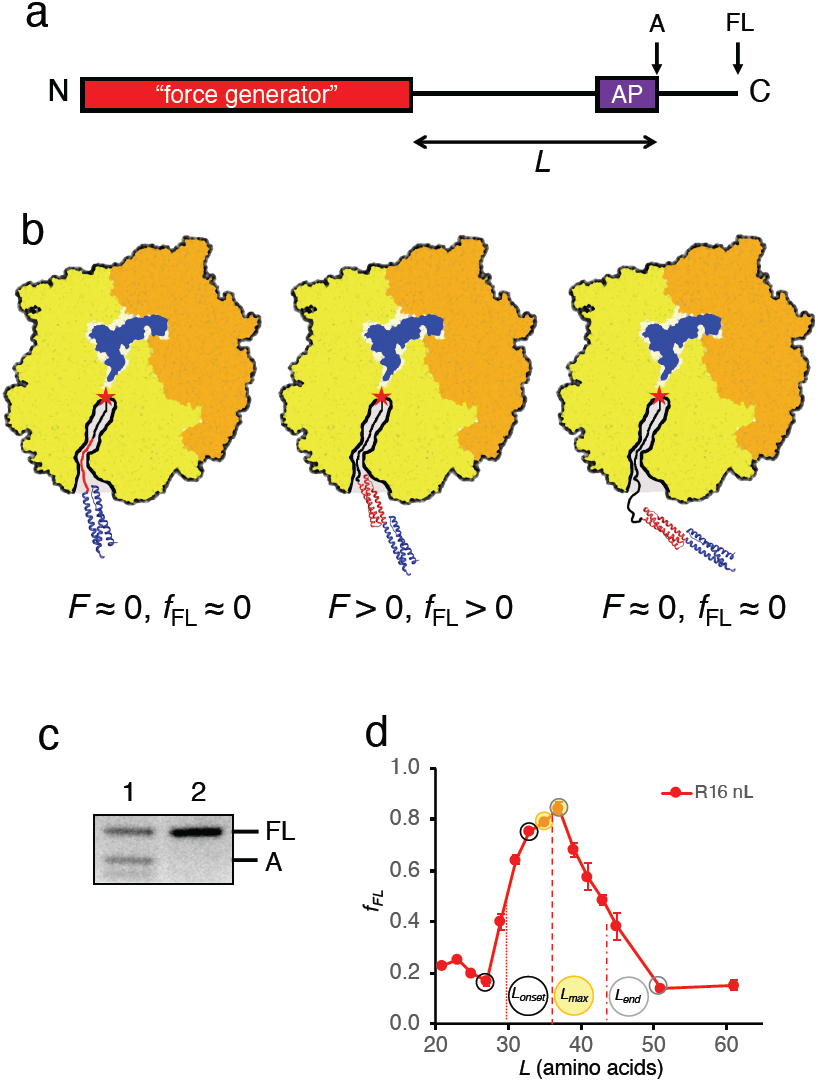
Force profile analysis. (a) Schematic representation of a typical construct. A “force-generator,” which here is one or more spectrin domains, is connected by a linker sequence to the *E. coli* SecM arrest peptide (AP). Following the AP is an unrelated protein sequence from the *E. coli* LepB protein that is added to allow the arrested (A) and full-length (FL) forms to be separable by SDS-PAGE. (b) Schematic representation of cotranslationally arrested ribosome-nascent chain complexes with increasing linker lengths (left to right). Represented here is the R15R16 nL construct (colored as in Fig. 2a). Left: at short linker lengths, when the folding of the R16 domain (red) has not yet begun, the force acting on the nascent chain is minimal, and the arrest is efficient (*F* ≈ 0, *f*_*FL*_ ≈ 0). Middle: at linker lengths that allow the folding of the R16 domain, the force acting on the nascent chain is increased and the arrest is less efficient (*F* > 0, *f*_*FL*_ > 0). Right: at linker lengths that arrest the ribosome after the folding of the R16 domain has already occurred, the force acting on the nascent chain is again minimal (*F* ≈ 0, *f*_*FL*_ ≈ 0). (c) Determination of *f*_*FL*_ by SDS-PAGE. Lane 1 is the R15 nL39 construct and lane 2 is the R15 nL39 full-length control construct. The control construct has the final (critical) proline residue of the AP mutated to alanine, preventing arrest and producing only full-length protein. (d) A plot showing the calculation of *L*_*onset*_, *L*_*max*_, and *L*_*end*_. The data points used for the calculation are denoted by black, yellow, and grey circles, respectively. The interpolated midpoint between each pair of selected point gives the *L*_*onset*_ (vertical dotted line), *L*_*max*_ (dashed line), and *L*_*end*_ (dash-dot line) values.

The cotranslational folding process can hence be followed by measuring the amount of full-length and arrested protein product at each linker length, Fig. 1c. Folding transitions will appear as peaks in a “force profile” plot of the fraction full-length protein (*f*_*FL*_) vs. linker length (*L)*, Fig. 1d, and can be described by the peak amplitude and the linker lengths that define the onset (*L*_*onset*_), maximum (*L*_*max*_), and end (*L*_*end*_) of the peak, as shown.

### Cotranslational folding of tandem spectrin domains

In the intact spectrin protein, the C-terminal α-helix in one domain is continuous with the N-terminal α-helix in the next domain, Fig. 2a. Consecutive domains are thus intimately connected to each other, and can influence each other’s thermodynamic stability (24, 27, 31, 42). In order to better understand the cotranslational folding of spectrin, we decided to systematically evaluate the effects of up- and downstream domains on the force profiles of the R15 and R16 spectrin domains by analyzing the two- and three-domain combinations shown in Fig. 2b. For each domain combination, a force profile was recorded by generating a series of constructs where the 17-residue SecM(*Ec*) AP (FSTPVWISQAQGIRAGP) from the *E. coli* SecM protein was placed at different linker lengths *L* from the C-terminal end of the R15 or R16 domain, translating each construct for 20 minutes in an *in vitro* translation system (43) in the continuous presence of [^35^S]-methionine, separating the translation products by SDS-PAGE, and calculating *f*_*FL*_ from the intensities of the bands corresponding to the full-length and arrested forms of the protein. The full amino acid sequences of all constructs are given in Supplementary Table S1.

**Figure 2.**
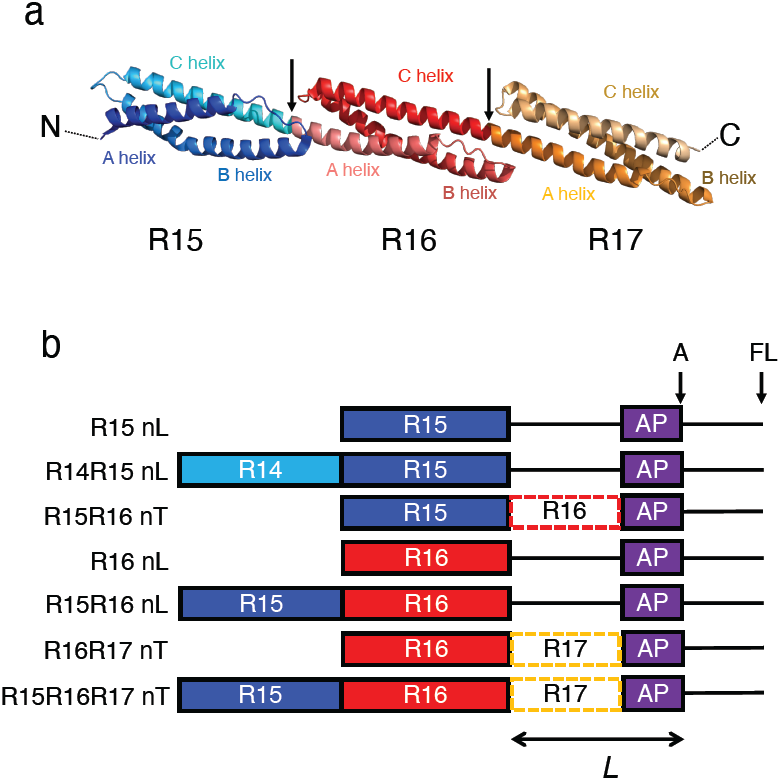
Construct design. (a) Structure of the R15 (blue), R16 (red), and R17 (orange) domains of chicken brain α-spectrin (PDB:1U4Q) (24). Arrows indicate the domain boundaries and the A, B, and C helices of each domain are labelled. (b) Constructs analyzed in this study. A solid black line represents sequence derived from the LepB protein and dashed boxes represent truncated spectrin sequences. The ends of the arrested (A) and full-length (FL) forms are indicated.

In order to distinguish which spectrin repeat is being assayed in the tandem constructs, we use the following naming convention. Constructs named “nL” (for **L**epB linker) are used to follow the folding of most C-terminal spectrin repeat (i.e, R15R16 nL follows the folding of R16) and constructs named “nT” (for **T**runcated spectrin linker) follow the folding of the penultimate spectrin repeat (i.e. R15R16 nT follows the folding of R15).

The results are shown in Fig. 3a-c. These include previous measurements collected for R16 nL and R15R16 nL (9) along with additional replicates and newly collected data for R15 nL. To facilitate comparison of the results, the *L*_onset_, *L*_max_, and *L*_end_ values extracted from each force profile are summarized in Supplementary Fig. S1; note that *L*_max_ values for force profiles that reach saturation (*f*_*FL*_ ≈ 1) cannot be accurately determined. As seen in Fig. 3b, the cotranslational folding of the R15 domain is unaffected by the presence of the upstream R14 domain (dark blue and light blue curves), while the presence of the N-terminal part of the downstream R16 domain induces a reduction in *L*_*onset*_ from 36 to 34 residues and a marked increase in the amplitude of the peak (compare the dark blue and purple curves). Thus, the onset of folding of R15 is sensitive to the presence of the early parts of the N-terminal α-helix of the downstream R16 domain, but is not affected by the presence of the upstream R14 domain.

**Figure 3.**
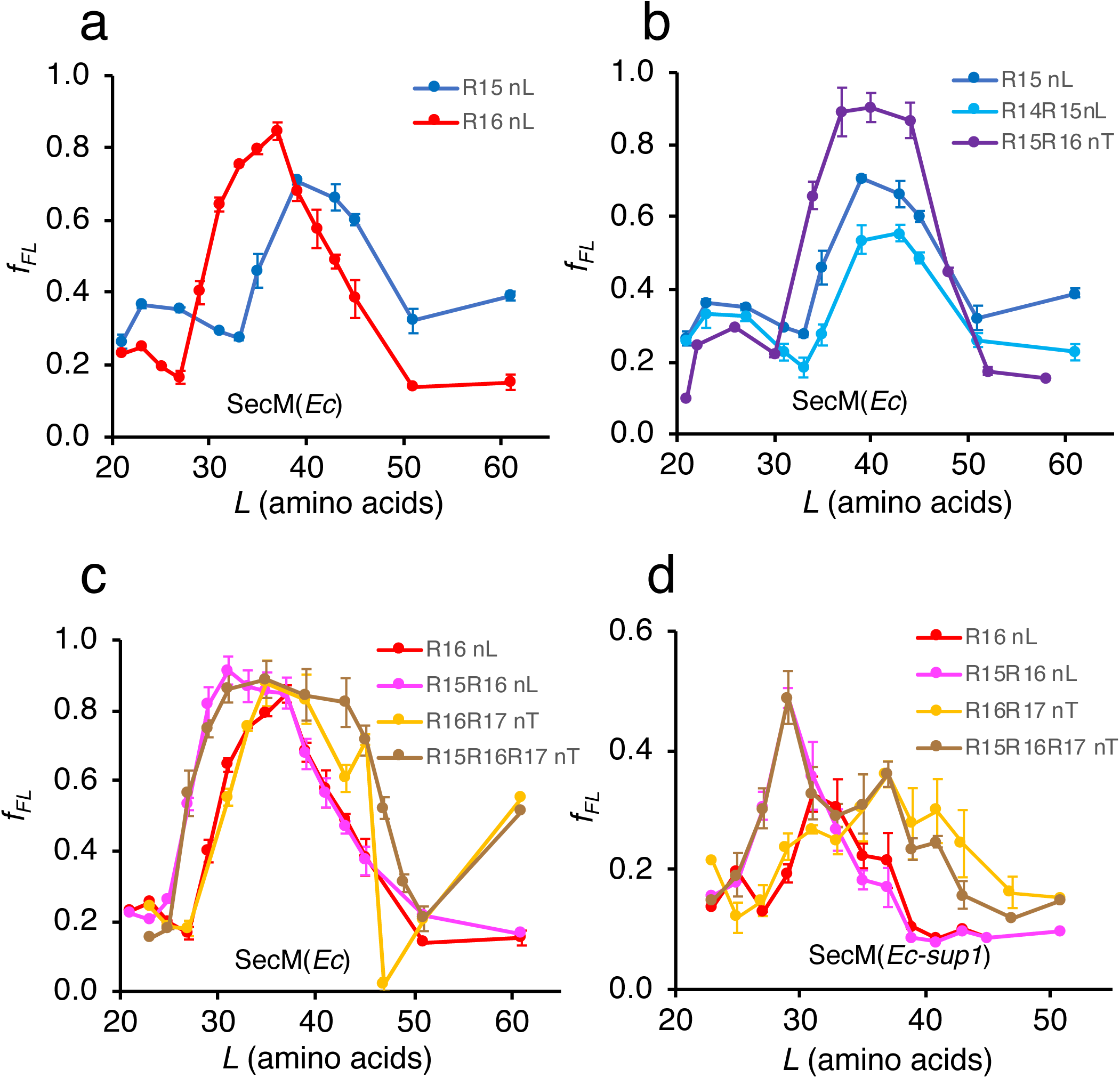
Cotranslational folding of spectrin domains. (a) Force profiles obtained with the SecM(*Ec*) AP for the R15 nL and R16 nL constructs. *f*_*FL*_ is the fraction full-length protein produced and *L* is linker length (*i.e*., the number of amino acid residues between the C-terminal end of the spectrin domain and the C-terminal Pro residue in the AP). Data for R16 nL constructs from (9) along with additional replicates are presented. (b) Force profiles obtained with the SecM(*Ec*) AP showing the influence of R14 and R16 on the folding of R15. (c) Force profiles obtained with the SecM(*Ec*) AP showing the influence of R15 and R17 on the folding of R16. Data for R15R16 nL from (9) along with additional replicates are presented. The reason for the abnormally low *f*_FL_ value for the R16R17 nT construct at *L* = 47 is unknown. (d) Force profiles obtained with the SecM(*Ec-sup1*) AP showing the influence of R15 and R17 on the folding of R16. Note that the R15R16 nL and the R15R16R17 nT profiles overlap perfectly in the interval *L* = 23-29 residues.

For the R16 domain, the effects of the neighboring R15 and R17 domains are more dramatic, Fig. 3c. The presence of the upstream R15 domain leads to a reduction in *L*_*onset*_ from 30 to 28 residues (compare the red and magenta curves), as shown before (9). The presence of the downstream R17 domain does not appreciably affect *L*_*onset*_ but leads to an increase in amplitude and a shift to a higher *L*_*end*_ value (compare the red and orange curves). Finally, when flanked by both the R15 and R17 domains, the R16 folding transition starts at a lower *L*_*onset*_ and ends at a higher *L*_*end*_ than for the isolated R16 domain (compare the red and brown curves).

In order to more precisely define *L*_max_ for constructs were *f*_*FL*_ approaches 1, we substituted two amino acids in the relatively weak SecM(*Ec*) AP to create the stronger SecM(*Ec*-*sup1*) variant (FSTPVWISQA<u>PP</u>IRAGP) (44) in the R16 nL, R15R16 nL, R16R17 nT, and R15R16R17 nT constructs, Fig. 3d. Again, the presence of R15 reduces *L*_*onset*_ and *L*_*max*_ by ∼2 residues (note that the R15R16 nL and the R15R16R17 nT profiles overlap perfectly in the interval *L* = 23-31), and the presence of R17 increases *L*_*end*_ by 6-7 residues (see Supplementary Fig. S2). Notably, the bi-modal shape of the R15R16R17 nT profile with peaks at *L* = 29 and 37 residues is an almost perfect match to the sum of the R15R16 nL and R16R17 nT profiles, Supplementary Fig. S2f, suggesting that the R16 part folds first (stabilized by the R15 C helix, c.f., Fig. 2a), followed by a second folding event involving the R17 A helix (stabilized by the R16 C helix).

Interestingly, both R16 folding constructs that include parts of R17 (R15R16R17 nT and R16R17 nT) show an increase in *f*_*FL*_ at the longest linker length (*L* = 61 residues), Fig. 3c. At this point the entire R17 A helix, the loop, and the very beginning of R17 helix B have been translated (see Table S1). This increase in *f*_*FL*_ may thus signal an early interaction between helices A and B in R17.

As a control, we introduced mutations that are known to inhibit folding of R15 and R16 *in vitro* (45, 46) into the R15 nL, R15R16 nT, R16 nL, R16R17 nT, and R15R16R17 nT constructs, Fig. 4 and Supplementary Fig. S3. As expected, *f*_*FL*_ values for the “non-folding” R15(nf) nL and R16(nf) nL constructs were strongly reduced, but, surprisingly, *f*_*FL*_ values remained high for linker lengths between *L*_*max*_ and *L*_*end*_ for R15(nf)R16 nT, R16(nf)R17 nT, and R15R16(nf)R17 nT constructs. Since the difference between the nL and nT series of constructs is the presence of a C-terminal “linker” derived from the N-terminal part of R16 or R17 respectively (replacing unstructured segments from LepB), this suggested that the linker itself, perhaps together with C-terminal parts of the upstream spectrin domain, might fold in the nf-mutants.

**Figure 4.**
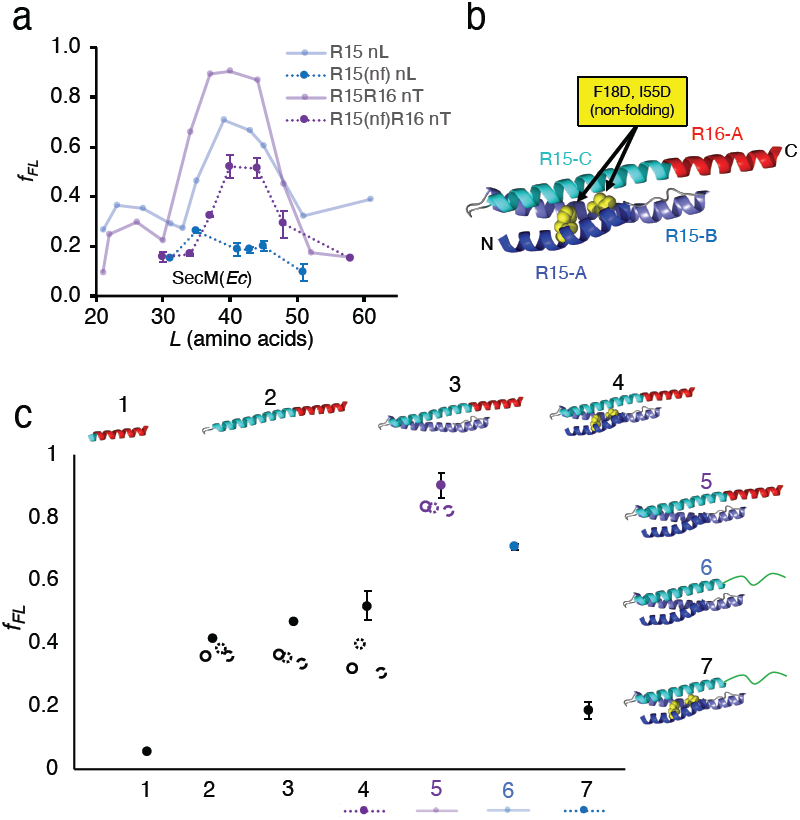
Linker effects in R15 folding. (a) Force profiles for R15 nL (blue, unbroken line) and R15R16 nT (purple, unbroken line) data from Fig. 3b compared with profiles for a R15 non-folding (nf) mutant (F18D + I55D) in R15(nf) nL constructs (blue, dashed line) and R15(nf)R16 nT constructs (purple, dashed line). (b) R15R16 nT spectrin structure (*L* = 40) with the A-C helices indicated and the destabilizing mutations F18D and I55D in the R15A and C helices shown as yellow spheres. (c) Graph showing *f*_*FL*_ values (*L* = 40) for constructs with increasingly long R15 parts fused to the N terminus of the 19-residue R16 nT linker (black circles, points 1, 2, 3, 4, and 5). R15 nL *L* = 39 (point 6) and R15(nf) nL *L* = 41 (point 7) are shown for comparison. Structural representations of each construct (spectrin helices are colored as in panel *b* and LepB linker sequences colored green) surround the plot. For points 1-4, the effect of helix-breaking insertions in the R16 A helix (Supplementary Table S1) in *f*_*FL*_ values are shown as open circles (Gly-Gly), open dotted circles (Pro), and open dashed circles (Gly-Ser-Gly-Ser).

To investigate this possibility, we started from the R15R16 nT, *L* = 40 construct (whose corresponding nf-mutant has a much higher *f*_*FL*_ value than the R15(nf) nL version, Fig. 4a) and successively removed helices A, B, and C from the R15 part (i.e. N-terminal truncations), Fig. 4b. We found that removing helix R15-A or both helices R15-A and R15-B decreased *f*_*FL*_, but only to 0.47 and 0.36, respectively, Fig. 4c (compare points #4, to #3, and #2). Only with the removal of all of R15, leaving only 19 amino acids of the R16 A helix, did *f*_*FL*_ decrease to baseline (point #1). Thus, the cotranslational formation of a continuous helical structure encompassing parts of the R15 C and R16 A helices appears to cause an increase in *f*_*FL*_ at *L* = 40; indeed, it is known from previous work that helices can form cotranslationally in the exit tunnel (47). The introduction of helix-breaking residues into the middle of the R16 A helix (see Table S1) in the R15-ABC/R16-A (point #4) and R15-BC/R16-A (point #3) constructs decreased the *f*_*FL*_ to the same level as R15-C/R16-A (point #2), Fig. 4c (open circles). Likewise, comparison of constructs where the R16 A helix in the R15R16 nT and R15(nf)R16 nT, at *L* = 40 was replaced by a non-helical segment from LepB at a comparable length (*L* = 39 and *L* = 41, respectively) led to reductions in *f*_*FL*_ (compare points #5 & 6 and #4 & 7). Similar behavior was also observed for the R16R17 nT, *L* = 43 constructs, Supplementary Fig. S3, implying that the R17 A helix forms a cotranslational folding intermediate together with the R16 C helix, as suggested above.

We conclude that the cotranslational folding of the R15 domain is affected by its downstream but not by its upstream neighbor domain, and that folding of R16 is affected both by its upstream and downstream neighbors. Different spectrin domains thus not only fold via different folding mechanisms, but their cotranslational folding is also differently impacted by their up- and downstream neighbor domains, a previously unappreciated feature of spectrin folding. We attribute the contribution of the downstream neighbor to the formation of helical structure encompassing helix C from the upstream domain and helix A from the downstream domain.

### A quantitative relation between f_FL_ and pulling force

In all FPA studies published to date, the quantitative relation between the calculated *f*_*FL*_ values and the underlying pulling force acting on the nascent chain has remained undefined (although attempts have been made to derive it from simulations or other kinds of theoretical modeling (11, 39)). Using the PURE translation system, Goldman *et al.* (6) showed that the interactions between the SecM(*Ec*) AP and the ribosome exit tunnel can be disrupted by a mechanical force applied through optical tweezers, and that the rate by which the translational stall induced by the SecM(*Ec*) AP is released, *k*_R_, increases in step with the amount of force, *F*, applied to the nascent chain, in a way that can be approximated by the Bell model (35) for force-induced rupture:

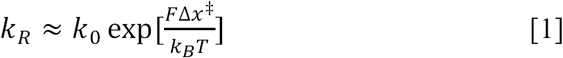

where *k*_0_ = 3 × 10^−4^ s^−1^ (95% confidence interval (CI): 0.5 × 10^−4^ s^−1^, 20 × 10^−4^ s^−1^) is the release rate at zero pulling force, and Δ*x*^‡^ = 0.4 nm (95% CI: 0.1 nm, 0.8 nm) is the distance to the transition state.

The work by Goldman *et al*. (6) suggested to us that approximating *k*_*R*_ with the rate of conversion of the arrested form of a given construct to the full-length form as measured in a pulse-chase experiment would allow us to estimate the corresponding pulling force using the relation in Eq. [1] between *F* and *k*_*R*_. To this end, we did pulse-chase experiments on a range of spectrin R16 and ADR1a (7) constructs that have *f*_*FL*_ values between 0.2 and 0.9, and fit the pulse-chase data to a first-order kinetic equation, Supplementary Fig. S4.

To derive an analytical relation between *f*_*FL*_ and *F*, we reasoned that if *f*_*FL*_ were measured at a single delay time (*Δt*) in a pulse-chase experiment, rather than under the standard continuous-labeling experimental conditions, the release rate (and hence *F*, according to Eq. [1]) could be estimated from:

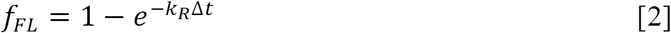

Since in the standard experiment translation can be initiated at any time during the 20 min. labelling period, the delay time *Δt* varies from one ribosome-nascent chain complex to another. Although many factors could in principle contribute to the distribution of delay times, it turns out empirically that using an average delay time *Δt* = 550 s, approximately equal to half of the total incubation time, describes remarkably well the relation between the standard *f*_*FL*_ values (20 min. continuous [^35^S]-Met labeling) and the release rates measured by pulse-chase experiments, Supplementary Fig. S5.

Combining Eqs. [1] and [2], one can solve for the pulling force *F*:

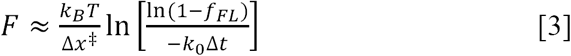

To obtain good estimates of the Bell parameters *k*_0_, Δ*x*^‡^ we started from the values obtained in (6) and performed a local optimization by requiring that *k*_*0*_ (the release rate at zero force) is close to *k*_*R*_ measured for the “zero-force” construct R16 nL27, Fig. 5a, and that the titin I27, spectrin R16, and ADR1a force profiles in (12) are well reproduced by the molecular dynamics simulation also described in (12). These conditions are both fulfilled by setting *k*_0_ = 3 × 10^−4^ s^−1^ (*i.e*., the same value as in (6) and equal to *k*_*R*_ for R16 nL27, Fig. 5a) and *x*^‡^ = 0.65 nm (well inside the 95% CI from (6)), Supplementary Fig. S6. With these parameter values and Δ*t* = 550 s, Eq. [3] nicely captures the relation between the force estimated from the measured *k*_*R*_ and standard *f*_*FL*_ values, Fig. 5b, and can hence be used to predict *F* from *f*_*FL*_.

**Figure 5.**
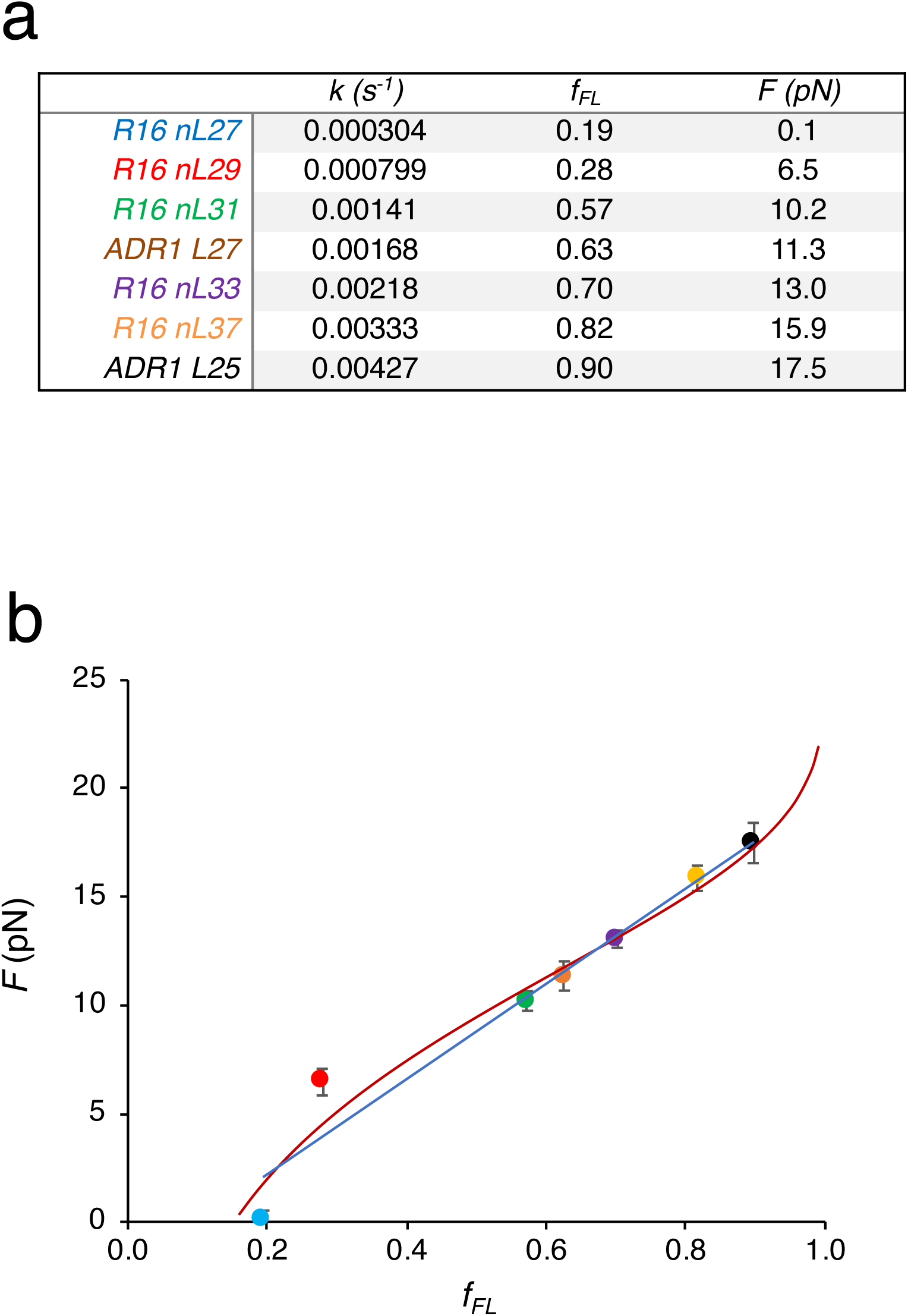
Release rates and estimation of pulling forces. (a) The rate of release (*k*_*R*_) obtained from pulse-chase experiments (see Supplementary Fig. S4 and Supplementary Table S2), the fraction full-length protein (*f*_*FL*_) measured under standard experimental conditions (20 min. incubation in PURExpress in the continuous presence of [^35^S] Met), and the pulling force *F* calculated using Eq. [1] (*k*_0_ = 3.0 × 10^−4^ s^−1^, Δ*x*^‡^ = 0.65 nm). The constructs are from Kudva *et al.* (12), and are colored to match those in panel *b* and Supplementary Figs. S4 and S5. (b) *F* values calculated from Eq. [1] plotted against the standard *f*_FL_ values, with constructs colored as in panel *a*. The least-squares fit line is indicated by the blue line, and the analytic relation Eq. [3] between *F* and *f*_*FL*_, assuming an average delay time Δ*t* = 550 s (approximately equal to half the standard incubation time), is shown as a red curve.

Over the interval 0.2 < *f*_*FL*_ < 0.9, the experimental data is also well approximated by the simple linear fit *F* = 22 *f*_*FL*_ – 2.2 (blue line). Obviously, Eq. [3] holds only for the 17-residue SecM(*Ec*) AP used here, and not for other APs of different stalling strengths (39). Presumably, Eq. [3] may be applied also in other contexts where the SecM AP is used to measure pulling forces, such as during membrane protein synthesis or translocation of charged residues across energized membranes (38, 39), although the parameters would first need to be verified to hold for *in vivo* experiments. We note that another approach to determining Bell parameters under a given set of conditions would be to repeat the FPA experiments using multiple APs with different resistance to force. A global fit of *f*_*FL*_ obtained for each AP as a function of linker length *L* would allow simultaneous determination of the forces exerted by the protein (independent of AP) as well as AP-dependent Bell parameters. In Supplementary Fig. S7 we illustrate this approach using data for translocon-mediated transmembrane helix insertion into the inner membrane of *E. coli* from *in vivo* AP measurements.

The force *F* estimated from pulse-chase measurements or *f*_*FL*_ values represents the constant force that would have the same effect on the escape rate from arrest as the combined effect of the forces from both folded and unfolded states. We can compare this inferred force with the ensemble average force calculated from MD simulations, ⟨*F*_*MD*_⟩ = *p*_*U*_*F*_*U*_ + *p*_*F*_*F*_*F*_, where *p*_*U*(*F*)_ and *F*_*U*(*F*)_ are, respectively, the population of, and force exerted by, the unfolded (folded) state in the simulation. In previous work (12), we calculated both *f*_*FL*_ as well as ⟨*F*_*MD*_⟩ directly from MD simulations. In Supplementary Fig. S8a, we show the relation between ⟨*F*_*MD*_⟩ and *f*_*FL*_ from our previous simulations (12), which is in good agreement with the relation between *F* and *f*_*FL*_ calculated from Eq. [3]. This agreement also confirms that the pulling forces are small enough that the force *F* calculated from simulated *f*_*FL*_ values using Eq. [3] is close to the ensemble-average force ⟨*F*_*MD*_⟩ determined by MD simulations; the two forces are not equal in general because of the non-linear relation between release rate and force, such that states exerting a larger force (e.g. the folded state) contribute disproportionately to the average release rate and hence to *F* (a direct comparison between ⟨*F*_*MD*_⟩ from the MD simulations and *F* calculated from Eq. [3] is given in Supplementary Fig. S8b).

We note that the forces exerted by a protein folding as it exits the ribosome are qualitatively different from the tensile forces such as those exerted when proteins fold in AFM experiments (as have been performed on spectrin before (48)). Therefore, the force magnitudes probed by the two experiments cannot be directly compared, despite their apparent resemblance. For example, even a small force exerted on the termini of an unfolded protein by an AFM can massively slow the refolding rate because of the large distance the protein must contract against this force in order to fold. The same force magnitude would have a much smaller effect on the refolding rate of a protein attached at one end to the ribosome (11).

## Discussion

α-spectrin contains over 20 repeat domains and is produced as a single, long polypeptide. Previously, we observed that the R16 domain starts to fold while a part of its C-terminal α-helix is still in the ribosome exit tunnel (9). We now find that the cotranslational folding of R16 is affected both by the presence of its upstream R15 domain and when the N-terminal part of the following R17 domain is present in the exit tunnel. In contrast, folding of the R15 domain is not affected by the upstream R14 domain, but starts at shorter linker lengths when the N-terminal end of the following R16 domain is present.

*In vitro*, R16 is stabilized by ∼1.7 kcal/mol by the presence of R15 (49), meaning that, during cotranslational folding, R16 can “sacrifice” a few interactions in the folding nucleus and start to fold at a shorter linker length in the R15R16 nL constructs (9).

For R15, the situation is a bit different because, in contrast to R16, its folding nucleus involves residues located at the C-terminal end of helix C (45). The presence of the upstream R14 domain thus may have little impact on *L*_*onset*_, because the whole R15 domain anyway must have emerged from the exit tunnel before folding can start. On the other hand, the R15 C helix is stabilized by the presence of the early parts of the R16 A helix, possibly allowing the folding nucleus to form at a shorter linker length and reducing *L*_*onset*_ in the R15R16 nT constructs.

The picture that emerges is that the spectrin domains fold one after the other as they emerge from the exit tunnel, but not completely independently of one another. As seen for the R16 domain, the presence of an already folded N-terminal upstream neighbor domain not only can increase the thermodynamic stability of the folded state of an emerging domain, but can also allow the emerging domain to start folding while still partly buried in the exit tunnel.

Likewise, when the C-terminal α-helix in the emerging domain is extended by a part of the N-terminal α-helix in the following domain, the folding transition can persist to longer linker lengths (as seen for R16) or start at shorter linker lengths (as seen for R15). Finally, the increase in *f*_*FL*_ seen for the R16R17 nT and R15R16R17 nT constructs at *L* = 61 residues (Fig. 3c) is suggestive of an early folding intermediate in R17. Further work is required to investigate this, but it may be possible that the regular helical structure of the spectrin repeats drives a nearly-continuous folding reaction, punctuated by periods when the nascent chain is lengthened but no residues are added to the folded part emerging from the ribosome. In this scenario, at most a short stretch of unfolded polypeptide would be exposed outside the ribosome at any one time, which could minimize the need for protection of the nascent chain by chaperones.

Finally, a pulse-chase analysis has allowed us to measure the release rate *k*_*R*_ from the stalled state for different R16 and ADR1a constructs, making it possible to estimate the magnitude of the pulling forces exerted on the AP for different linker lengths, Fig. 5a. This in turn makes it possible to derive expressions for how the magnitude pulling force *F* depends on *f*_*FL*_ (as obtained from our standard continuous-labeling experimental protocol), Eqs. [2] and [3]. In general, we find that the folding of protein domains such as spectrin R16 and ADR1a can generate a maximal force of 15-20 pN on the AP, Fig. 5, in line with theoretical estimates based on MD simulations (40, 50).

## Materials and Methods

All the enzymes used for molecular biology were obtained from New England Biolabs. The PUREfrex *in vitro* transcription-translation system was produced by Eurogentec and purchased via BioNordika. The PURExpress *in vitro* transcription-translation system was purchased from NEB. GeneJet PCR clean-up kit was purchased from Thermo-Fisher Scientific. Precast NuPAGE gels and running buffers were purchased from Invitrogen. Homemade gels were cast using acrylamide-bisacrylamide mix, Tris, and glycine from VWR. Filter paper for gel drying was from Whatman. All other chemicals were purchased from Sigma.

### Cloning

The cDNAs for spectrin R15 and R16 domains were kindly provided by Dr. Jane Clarke and the spectrin R17 and R14 domains were ordered as DNA fragments from Eurofins Genomics GmbH. All protein sequences used in this publication are presented in Supplementary Table S1. We used two different construct types: those with an unrelated linker sequence, labelled nL, and those with a truncated spectrin linker sequence, labelled nT (Fig. 2b). For nL constructs, the respective spectrin domain or domains were cloned into the pET19b plasmid upstream from a Ser-Gly-Ser-Gly sequence attached to a linker sequence derived from the P2 domain of leader peptidase (LepB) followed by the 17-residue AP from the *E. coli* SecM protein (FSTPVWISQAQGIRAGP) herein referred to as SecM(*Ec*), and a further 23 amino acids derived from LepB. The shortest constructs (nL/nT 21) include only the SGSG sequence fused to the SecM(*Ec*) AP. For nT constructs, the respective spectrin domains were cloned into the same pET19b plasmid without a LepB linker sequence, but including the SGSG sequence immediately before the SecM(*Ec*) AP. The most C-terminal spectrin domain was then sequentially truncated via partially overlapping inverse PCR. For some constructs, a full-length (FL) control was created by changing the critical Pro at the C-terminal end of the SecM AP to Ala, thereby abolishing arrest (51) and yielding only full-length protein. The SecM(*Ec-sup1*) variants were created by site-directed mutagenesis of the constructs described above. N-terminal truncations of R15 and R16 were carried out by partially overlapping inverse PCR and insertion of the helix breaking insertions was carried out by site-directed mutagenesis where the inserted residues were included in the primer sequence. All sequences were confirmed by DNA sequencing (Eurofins Genomics GmbH).

### In vitro expression for Force Profile Analysis (FPA)

Expression and analysis was carried out as described previously (9). Briefly, a linear DNA product is created from each construct plasmid by PCR using Q5 polymerase with forward and reverse primers that anneal to the T7 promoter and terminator regions, respectively. Following PCR clean-up (using the manufacturer’s instructions) the product is confirmed by agarose gel electrophoresis. *In vitro* transcription and translation is carried out in either the PUREfrex of PURExpress commercial systems (mixed according to the manufacturer’s recommendations). 1 µL of the PCR product and ∼10 µCi [^35^S]-methionine are mixed for a 10 µL PUREfrex reaction or 0.5 µL of PCR product and ∼5 µCi [^35^S]-methionine are mixed for a 5 µL PURExpress reaction, followed by incubation at 37°C for 20 min at 600 rpm shaking. Translation is halted by the addition of an equal volume of ice-cold 10% TCA, followed by incubation on ice for at least 30 min. Total protein is sedimented by centrifugation at 4°C for 5 min at 20,000x*g*. The supernatant is carefully removed and the pellet is resuspended in a suitable volume of 1X SDS-PAGE sample buffer (62.5 mM Tris-HCl pH 6.8, 10% glycerol, 2.5% SDS, 5% β-ME, 0.02% bromophenol blue, and 25 mM NaOH (NaOH is added to neutralize any remaining TCA) by shaking at 37°C and 1,000 rpm for at least 5 min. The prolyl-tRNA that remains attached due to SecM arrest is digested by the addition of 4 µg of RNAse I (2 µL of a 4 µg/µL solution), followed by incubation at 37°C and 600 rpm for 30 min.

### Quantifications

Following a brief centrifugation to remove any remaining insoluble material, the sample is loaded onto an appropriate SDS-PAGE gel (12% tris-glycine gels were used for two- and three-spectrin domain constructs and 16 or 18% tris-tricine gels were used for single-spectrin domains and the N-terminal truncations). Following electrophoresis, the gels are dried onto thick filter paper by heating under vacuum (BioRad Model 583 or Hoefer GD 2000), a radioactive molecular weight ladder included in the gel is visualized by spotting the filter paper with a ∼1:1,000 solution of [^35^S]-methionine in 1X SDS-PAGE sample buffer, and the gel is exposed to a phosphorimager screen for 12 to 72 hours depending on the strength of the signal. The screen is imaged using a Fujifilm FLA9000 (50 µm pixels), and densitometry analysis on the resultant raw image (TIFF format) file is carried out using FIJI (ImageJ) software. The densitometry values are quantified using our in-house EasyQuant software and the fraction full-length protein, 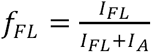, is calculated from the intensities of the full-length (*I*_*FL*_) and arrested (*I*_*A*_) bands. See Supplementary Fig. S9 for examples of gels. Independent replicate *in vitro* translation reactions were carried out for all spectrin constructs in the folding peaks, see Supplementary Table S2 (191 unique constructs and 405 independent data points). The majority of the data points in the folding peaks were collected in triplicate, with the remaining points collected in duplicate. A small number of data points outside of the folding peaks were collected as single measurements to save costs.

### Pulse-chase experiments

The rate at which translation recommences following the arrest of various constructs was measured for five R16 constructs and two ADR1 constructs (7) chosen to represent the range of *f*_*FL*_ values measured during typical experimental conditions. Pulse-chase experiments were carried out using the PURExpress system. 75 µL reactions were mixed according to the manufacturer’s instructions, with the addition of [^35^S]-methionine. Following a 5 min incubation at 37°C (pulse) an excess of unlabeled methionine was added and incubation at 37°C was continued. At discrete time points, a 10 µL aliquot of the reaction was removed and mixed with 15 µL of ice-cold 10% TCA and further processed as above. Three independent replicates were collected for each construct (see Supplementary Table S2). The rate calculated is for the conversion of arrested protein to full-length protein:

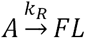

The rate of this irreversible conversion can be calculated using a 1^st^ order equation:

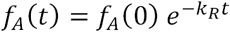

Where *f*_*A*_(t) is the fraction arrested protein at a given time, *t*, and *f*_*A*_(0) is the fraction arrested protein at *t* = 0. Prism 8 (GraphPad Software) was used to calculate the non-linear regression fit of the kinetic equation to each data set (R16 nL-27, -29, -31, -33 and -37, and ADR1 L25 and 27 (7)). The rates were used to calculate the force exerted by the folding domain on the arrested nascent chain as detailed in the main text.

### Optimization of the Bell parameters

The coarse-grained molecular dynamics (MD) simulations of cotranslational folding of R16 and ADR1a constructs in WT, ΔuL23, and ΔuL24 ribosomes used here were originally reported in (12). For each construct and each linker length, these simulations were used to calculate the ensemble average force ⟨*F*_*MD*_⟩ = *p*_*U*_*F*_*U*_ + *p*_*F*_*F*_*F*_, where *p*_*U*(*F*)_ and *F*_*U*(*F*)_ are, respectively, the population of, and force exerted by, the unfolded (folded) state in the simulation. In order to determine the Bell parameters *k*_0_ and Δ*x*^‡^ in Eq. [1] in the main text, we set *k*_0_ = 3 × 10^−4^ s^−1^ (*i.e*., the same value as estimated in (6), and equal to *k*_*R*_ for the “zero force” construct R16 nL27, Fig. 5a) and explored a range of Δ*x*^‡^ values around the approximate value Δ*x*^‡^ = 0.4 nm given in (6). For each Δ*x*^‡^ value, we calculated an average *k*_*R*_ over unfolded and folded states as *k*_*R*_ = *p*_*U*_*k*_0_ exp[*F*_*U*_ Δ*x*^‡^/*k*_*B*_*T*] + *p*_*F*_*k*_0_ exp[*F*_*F*_ Δ*x*^‡^/ *k*_*B*_*T*] and then obtained *f*_*FL*_ (setting Δ*t* = 550 s) from Eq. [2] in the main text. The simulated *f*_*FL*_ values (*i.e*, the simulated force profiles) obtained in this way were then compared with the experimental force profiles reported in (12) for the I27 and R16 domains expressed with WT, ΔuL23, and ΔuL24 ribosomes, and for the ADR1a domain expressed with WT and ΔuL24 ribosomes. The parameter values *k*_0_ = 3 × 10^−4^ s^−1^, Δ*x*^‡^ = 0.65 nm were found to give the best fit between the simulated and experimental force profiles (see Supplementary Fig. S7).

## Author contributions

GK, OBN, and GvH conceived the project. GK and OBN designed and performed the experiments. PT and RB performed the theoretical calculations. All authors contributed to writing the manuscript.

## Acknowledgements

This work was supported by grants from the Knut and Alice Wallenberg Foundation (2012.0282), the Swedish Cancer Society (15 0888), and the Swedish Research Council (621-2014-3713) to GvH, and from the Intramural Research program of the National Institute of Diabetes and Digestive and Kidney Diseases of the NIH to RBB, and from NIH (grant R35GM122543) to FPA. This work utilized the computational resources of the NIH HPC Biowulf cluster. (http://hpc.nih.gov). We thank Dr. Jane Clarke and Dr. Adrian Nickson for providing plasmids with spectrin DNA and Dr. Rickard Hedman for programming and maintenance of the EasyQuant software.

## Data Availability

The quantified raw data points are provided in Supplementary Table 2.

---

**Supplementary Figure S1.**
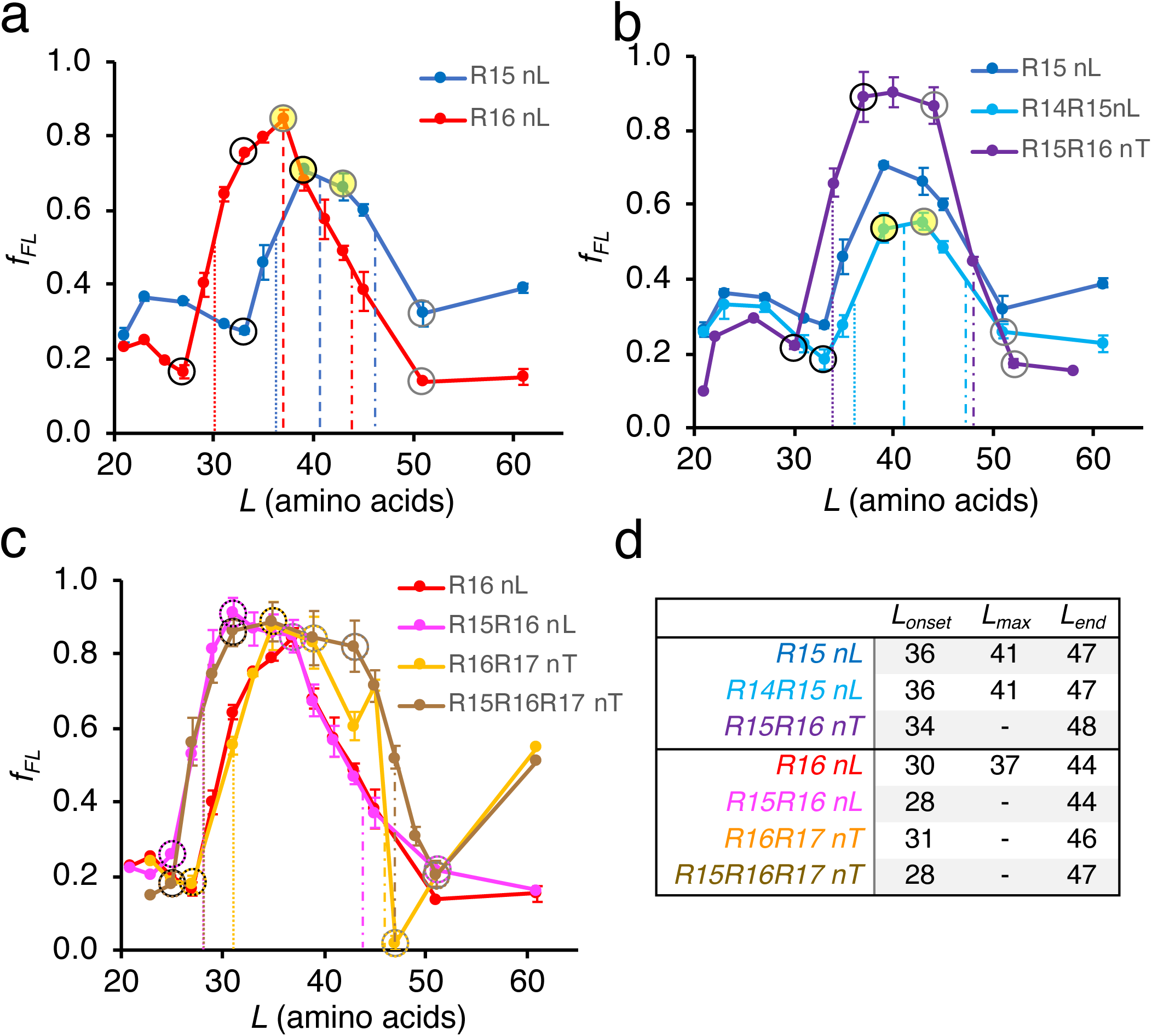
Calculation of *L*_*onset*_, *L*_*max*_, and *L*_*end*_ for all SecM(*Ec*) constructs. In panels a-c the data points used to calculate *L*_*onset*_ are indicated by black rings, *L*_*max*_ by filled yellow circles, and *L*_*end*_ by grey rings for each force-profile curve. For extra clarity, in panel c the rings are also decorated in the same color as the dataset. The *L* value at the midpoint between the selected data points is indicated by a dotted line (*L*_*onset*_), a dashed line (*L*_*max*_), or a dash-dot line (*L*_*end*_) in the same color as the respective dataset. (a) Force profiles for the R15 nL and R16 nL series of constructs. (b) Force profiles showing the influence of R14 and R16 on the folding of R15. (c) Force profiles showing the influence of R15 and R17 on the folding of R16. (d) Table of calculated *L*_*onset*_, *L*_*max*_, and *L*_*end*_ values for SecM(*Ec*) constructs. No *L*_*max*_ values were calculated for curves reaching saturation (*f*_*FL*_ ≈ 1).

**Supplementary Figure S2.**
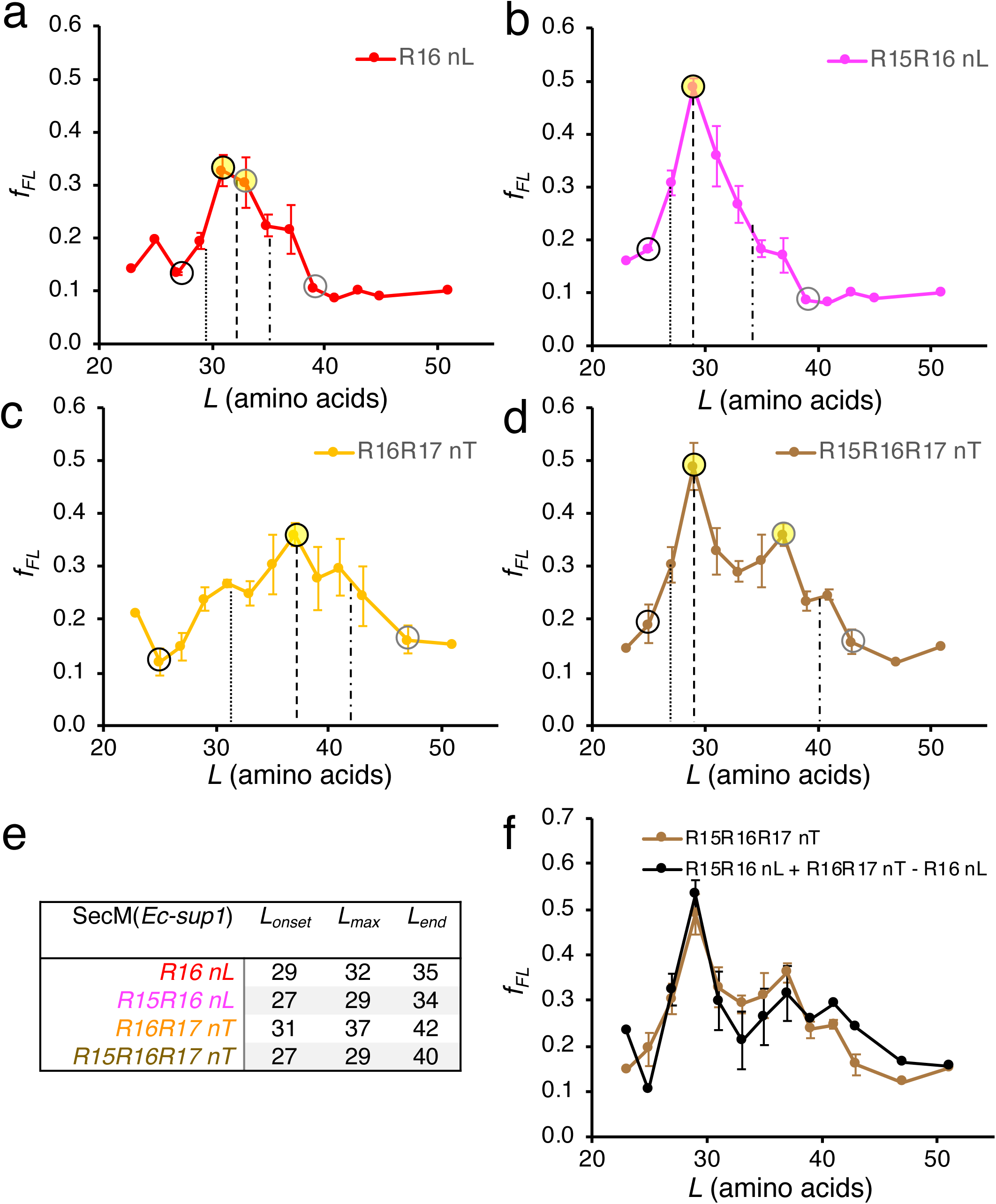
Calculation of *L*_*onset*_, *L*_*max*_, and *L*_*end*_ for all SecM(*Ec-sup1*) variants monitoring R16 folding (see Fig. 3d and Table S1). In panels a-c the data points used to calculate *L*_*onset*_ are indicated by black rings, *L*_*max*_ by filled yellow circles, and *L*_*end*_ by grey rings for each force-profile curve. The *L* value at the midpoint between the selected data points is indicated by a dotted line (*L*_*onset*_), a dashed line (*L*_*max*_), or a dash-dot line (*L*_*end*_). (a) Force profile for R16 nL. (b) Force profile for R15R16 nL. (c) Force profile for R16R17 nT. (d) Force profile for R15R16R17 nT. (e) Table summarizing the calculated values. (f) Sum of the R15R16 nL and R16R17 nT SecM(*Ec-sup1*) force profiles (blue) plotted with the R15R16R17 nT profile (brown). The summed profile was calculated by summing the individual profiles and subtracting the R16 nL profile to correct for double-counting of the R16 nL profile (see Supplementary Table S2).

**Supplementary Figure S3.**
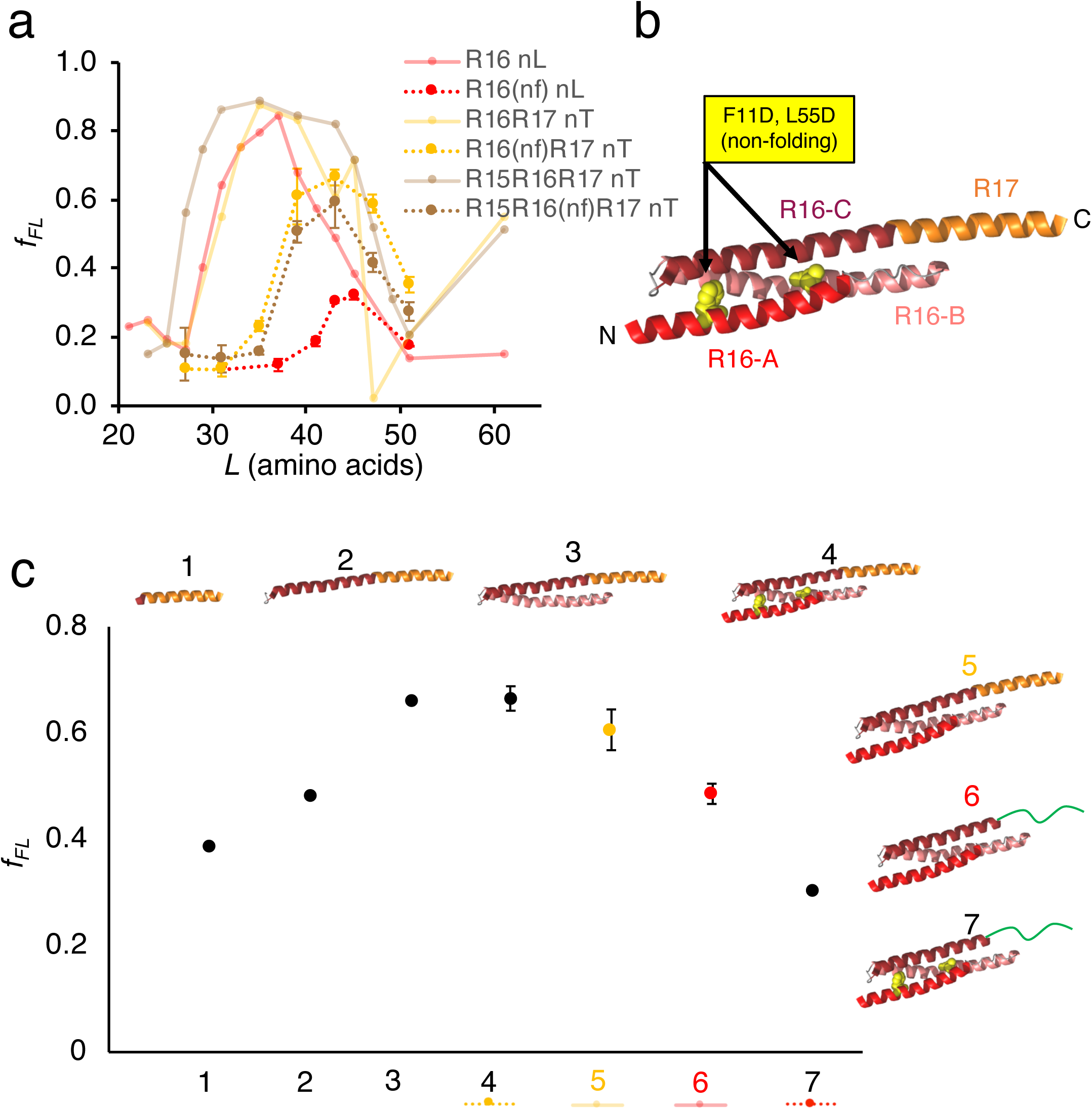
Investigation of the linker effect on R16 folding. (a) Reproduction of the R16 nL (red), R16R17 nT (orange), and R15R16R17 nT (brown) data from *Figure 3b* along with R16 point mutations that prevent folding (nf, F11D and L55D) in R16 nL (red, dashed line), R16R17 nT (orange, dashed line), and R15R16R17 nT (brown, dashed line). (b) R16R17 nT (*L* = 43) spectrin structure with helices R16-A, R16-B, R16-C, and R17-A labelled. (c) Graph showing the resultant *f*_*FL*_ at *L* = 43 with different N-terminal sequences fused to the R17 nT linker (black circles). A structural representation of each protein are shown for reference (spectrin helices are colored as in panel b and LepB sequences colored green).

**Supplementary Figure S4.**
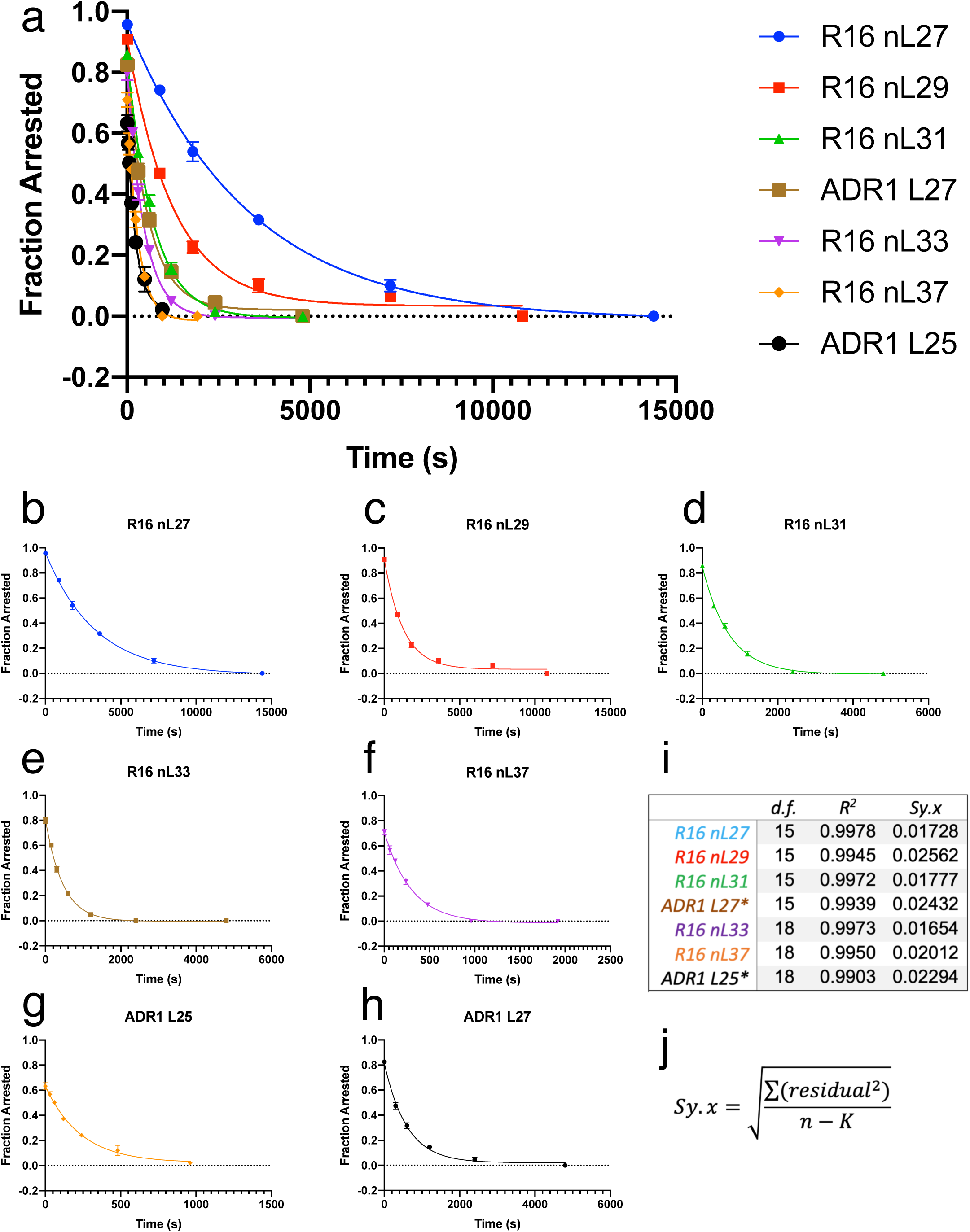
Determination of *k*_*R*_ for each construct. (a) Scaled overlay of the individual plots in panels b-h. (b)-(h) Plot of the decrease in fraction arrested protein (*f*_*A*_) over time after chasing a 5 min PURExpress translation of the indicated construct. The data were fit to the first order equation in the main text to determine the rate of release (*k*_*R*_) of arrested protein from the ribosome. (i) Summary of the fitness of the equation for each construct: degrees of freedom (*d.f.*), *R*^*2*^, and goodness-of-fit calculation (*Sy.x*). (j) The goodness-of-fit calculation provided by Prism 8 (Graphpad software) where *n* is the number of data points and *K* is the degrees of freedom.

**Supplementary Figure S5.**
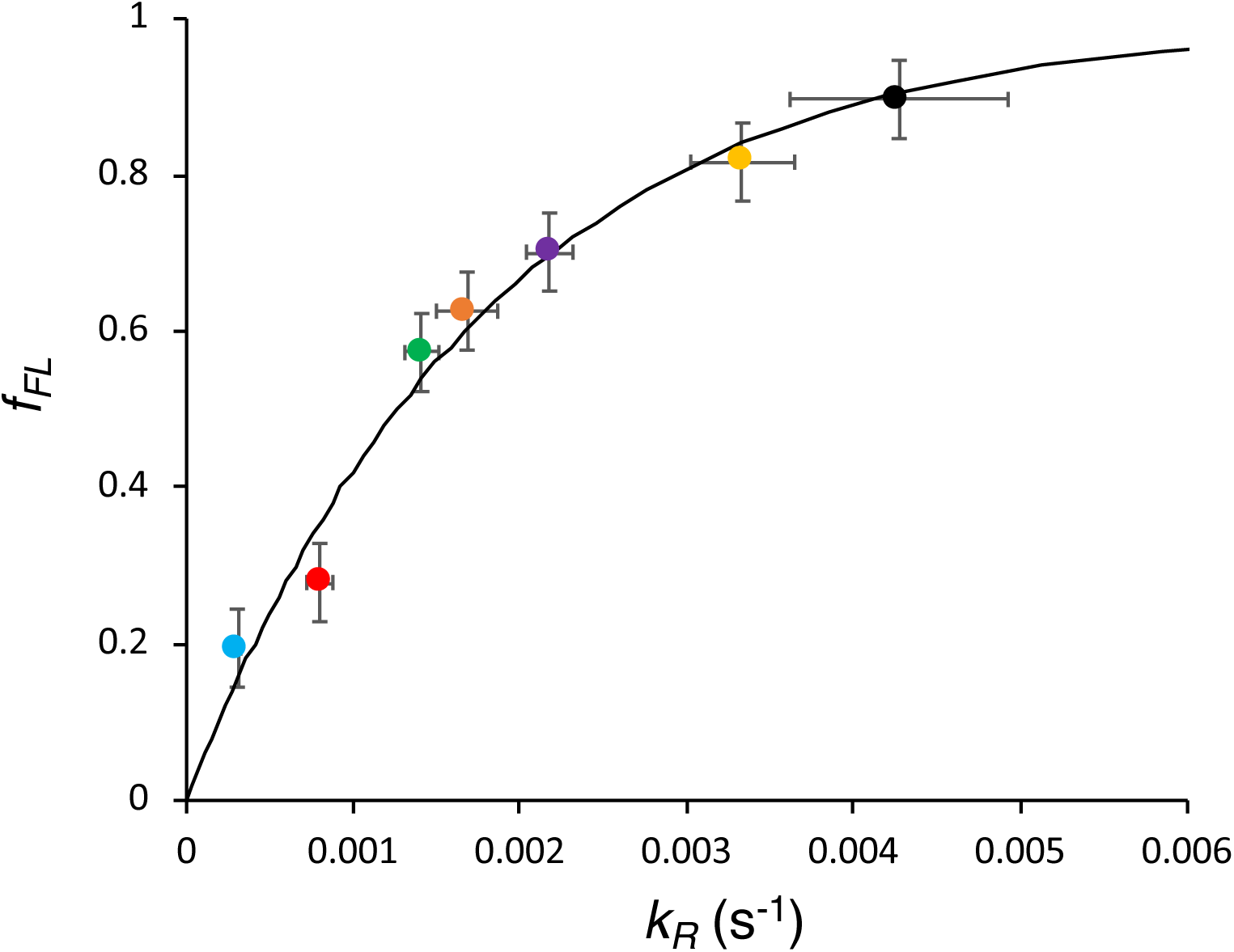
Empirical relation between standard *f*_*FL*_ values (20 min. continuous [^35^S]-Met labeling) and the release rate *k*_R_ measured in pulse-chase experiments. The data is fitted by the exponential *f*_*FL*_ = 1 − exp [−*k*_*R*_Δ*t*] (black curve) with Δ*t* = 550 s, i.e., approximately half of the full incubation time. Data points are colored as in Fig. 4. Error bars show 95% CI for *k*_*R*_ and the typical standard error (= 0.05) (Tian et al., *Proc Natl Acad Sci USA* 115, E11284 (2018)) for *f*_*FL*_.

**Supplementary Figure S6.**
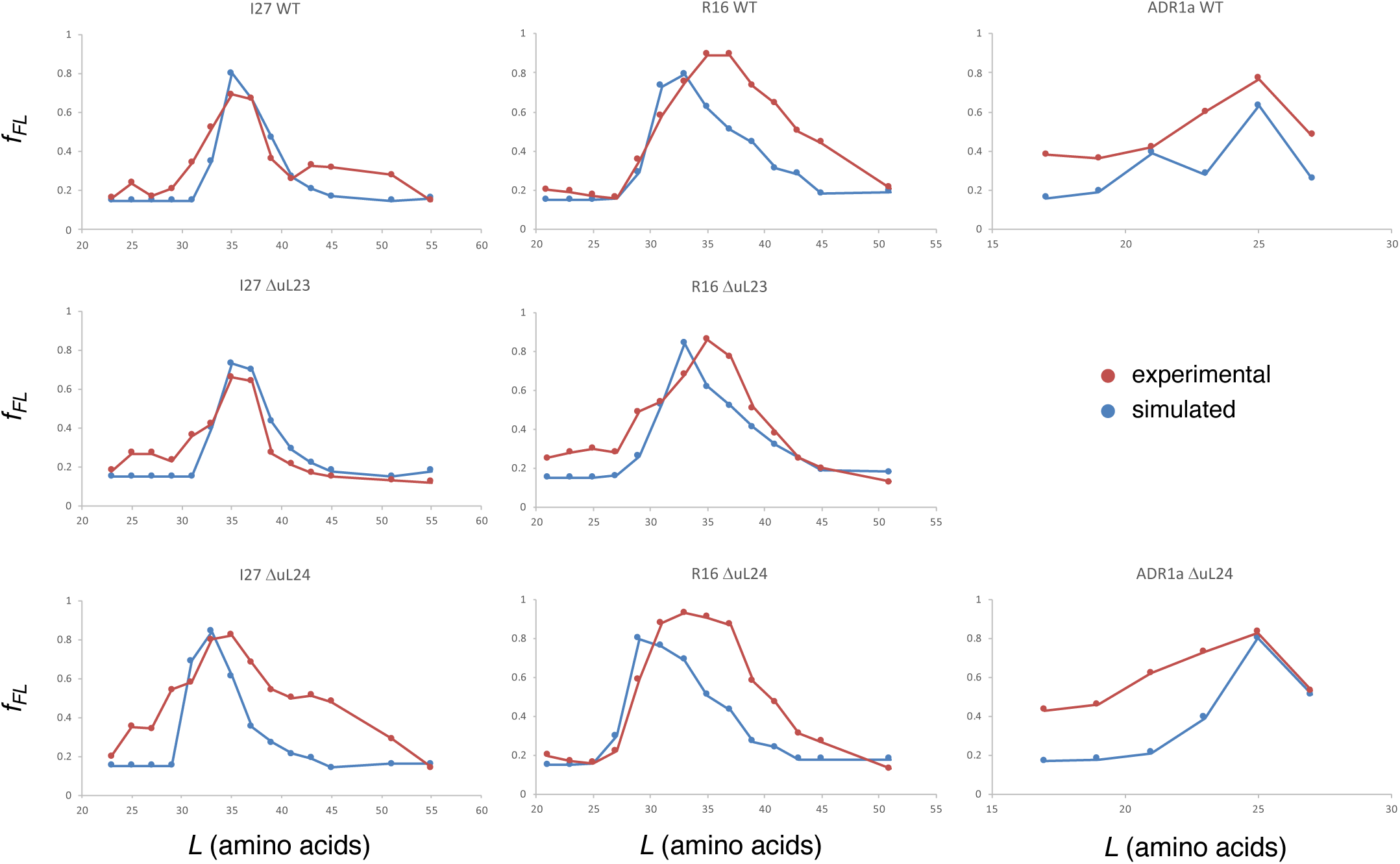
Simulated (blue) and experimental (dark red) force profiles for I27, R16, and ADR1a measured with wildtype (WT) and mutant (ΔuL23, ΔuL24) ribosomes (Kudva et al., *eLife* 7, 274191 (2018)). The simulated profiles were obtained as described in the *Materials and Methods* section, using the optimized Bell parameters *k*_0_ = 3 × 10^−4^ s^−1^ and *x*^‡^ = 0.65 nm. As discussed in (Kudva et al., *eLife* 7, 274191 (2018)), the simulations reproduce *L*_*onset*_, *L*_*max*_, and the maximal amplitude well for all the force profiles, but underestimate *L*_*end*_ for the I27 ΔuL24, R16 WT, and R16 ΔuL24 ribosomes. Since it is generally difficult to fit the simulated force profile to the experimental ADR1a ΔuL23 force profile described in (Kudva et al., *eLife* 7, 274191 (2018)), this was not included in the optimization of the Bell parameters.

**Supplementary Figure S7.**
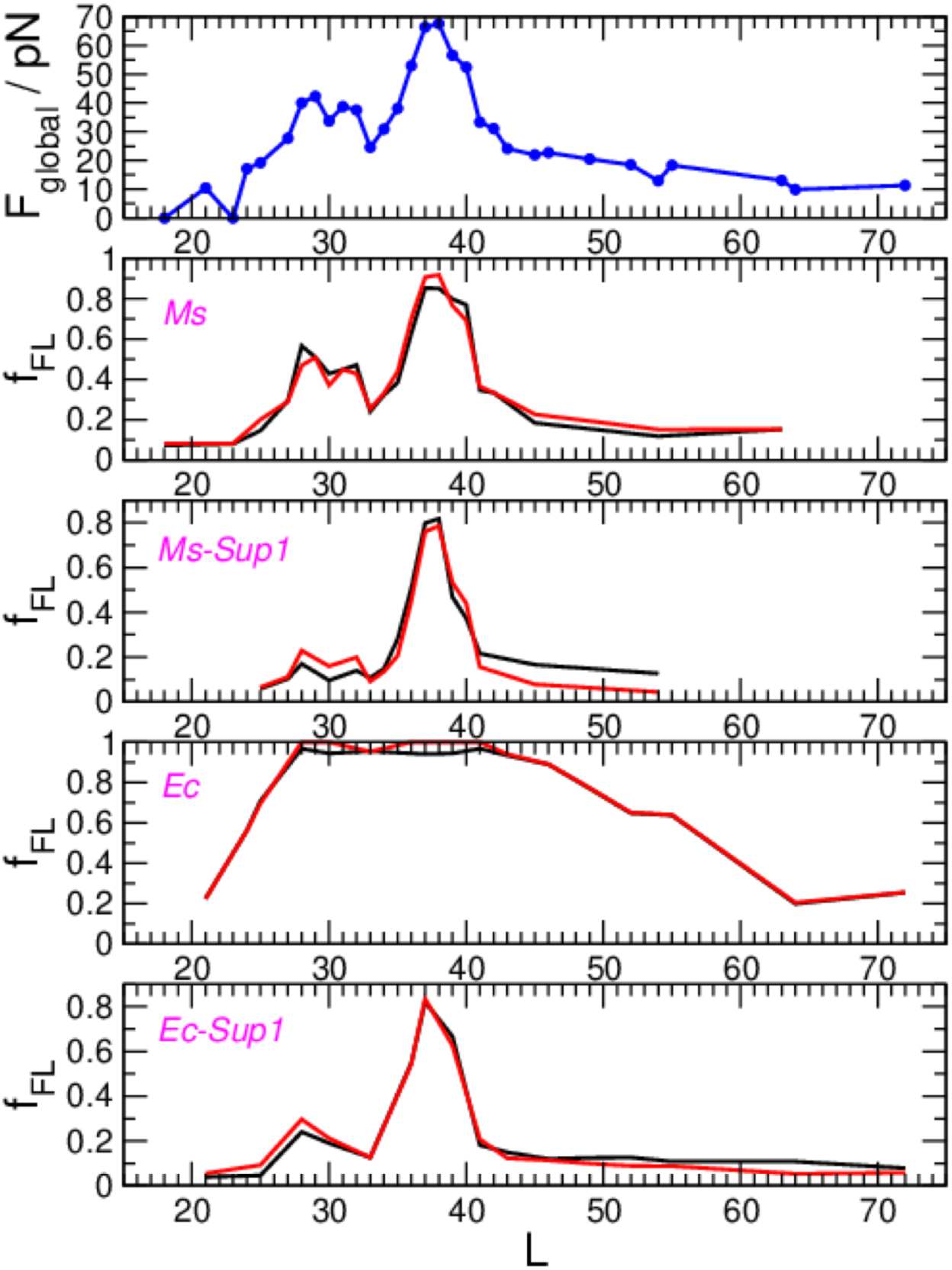
Global fit to the experimental data for translocon-mediated insertion of a transmembrane segment of composition [6L, 13 A] into the *E. coli* inner membrane from Ismail et al. (*Nat. Struc. Mol. Biol.* 19, 1018-1022 (2012)) to a common set of pulling forces. For each of the SecM arrest peptide (AP) variants *Ms, Ms-Sup1, Ec, Ec-Sup1* used in that work, an AP-specific set of Bell parameters *k*_0_, Δ*x*^‡^ is defined. In addition it is assumed that the force exerted on the AP depends only on the linker length *L*, i.e. *F*_*global*_*(L)*. The escape rate for each peptide and linker length can then be determined as 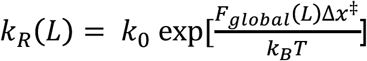, and hence the fraction full-length protein *f*_*FL*_ is obtained from *f*_*FL*_(*L*) = 1 − exp [− *k*_*R*_(*L*)*t*_*i*_], where *t*_*i*_ is the incubation time (120 s in these experiments). The AP-independent forces *F*_*global*_*(L)* and AP-dependent Bell parameters are globally fitted by minimizing the least-squares difference between experimental and calculated *f*_*FL*_ using a basin-hopping algorithm. The Bell parameters for the *Ec* AP were restrained to be similar to those found in Goldman et al (*Science*, 348, 457-460 (2015)). The best-fit global forces are shown in the top panel above, and the experimental (black curves) and calculated (red curves) *f*_*FL*_ data are shown in the lower panels for each AP. The optimal Bell parameters were found to be: *Ms: k*_0_ = 7.1×10^−4^s^−1^; *x*^‡^ = 0.21 nm; *Ms-Sup1: k*_0_ = 1.64×10^−4^s^−1^; *x*^‡^ = 0.27 nm; *Ec: k*_0_ = 3.3×10^−4^s^−1^; *x*^‡^ = 0.74 nm; *Ec-Sup1: k*_0_ = 2.4×10^−4^s^−1^; *x*^‡^ = 0.26 nm.

**Supplementary Figure S8.**
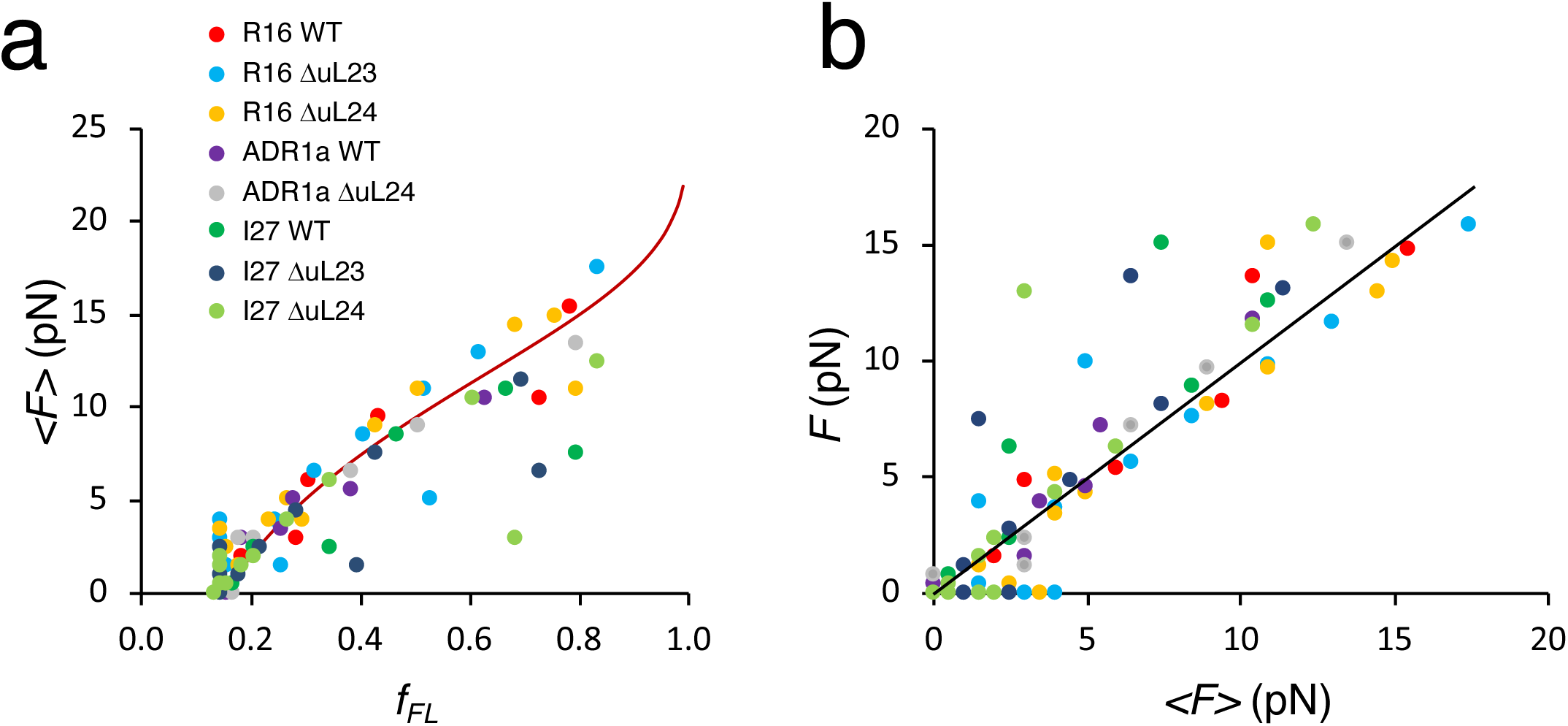
Force calculations based on coarse-grained molecular dynamics simulations of I27, R16, and ADR1a force profiles measured with wildtype (WT) and mutant (ΔuL23, ΔuL24) ribosomes (Kudva et al., *eLife* 7, 274191 (2018)). (a) Relation between the ensemble average force ⟨*F*_*MD*_⟩ and *f*_*FL*_ from simulation for the indicated constructs. The red curve is the same as in Fig. 4b, relating the force *F* from AP measurements to *f*_*FL*_. (b) Comparison of the ensemble average force ⟨*F*_*MD*_⟩ with the force *F* calculated from simulated *f*_*FL*_ values using equation [3]. The black line represents ⟨*F*_*MD*_⟩ The similarity of the forces indicates that they are small enough that ⟨*F*_*MD*_⟩ can be well approximated by *F*. In general *F* is expected to be slightly larger than ⟨*F*_*MD*_⟩.

**Supplementary Figure S9.**
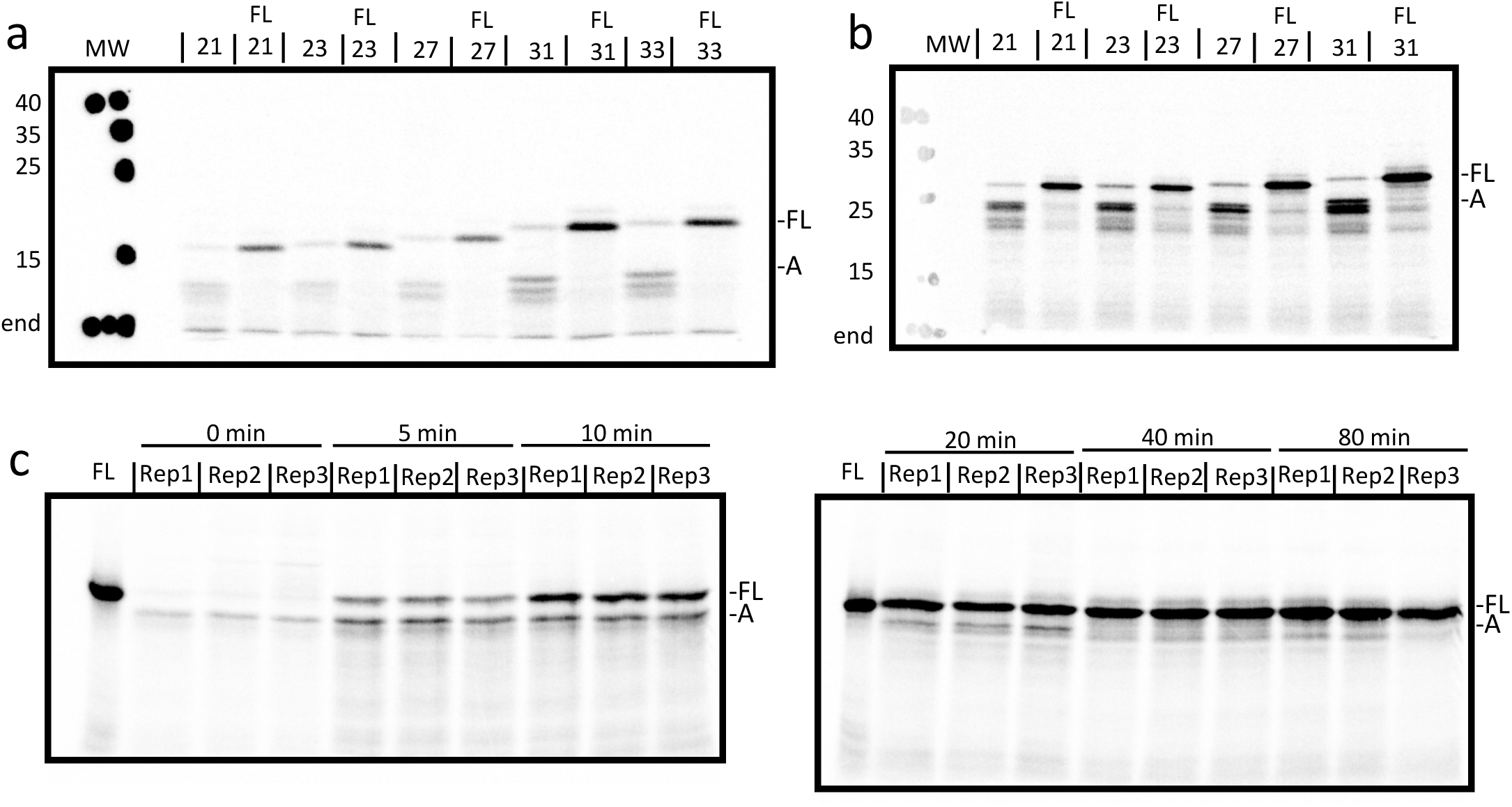
Examples of gels used for quantification of *f*_*FL*_. Above any lane FL indicates a full-length control construct (with an inactivated AP, see *Supplementary Table 1*). Location of full-length (FL) and arrested (A) products from the rightmost construct in each gel are indicated. (a) R15 nL. Construct *L* values are indicated above each lane and MW marker bands (kDa) are denoted as spots and labelled to the left of the gel. (b) R14R15 nL. Construct *L* values are indicated above each lane and MW marker bands (kDa) are denoted as spots and labelled to the left of the gel. (c) ADR1 L27 pulse-chase experiments. Triplicate samples (Rep1, Rep2, Rep3) at various time points (labelled in minutes) are above the lanes.

**Supplementary Table S1.**
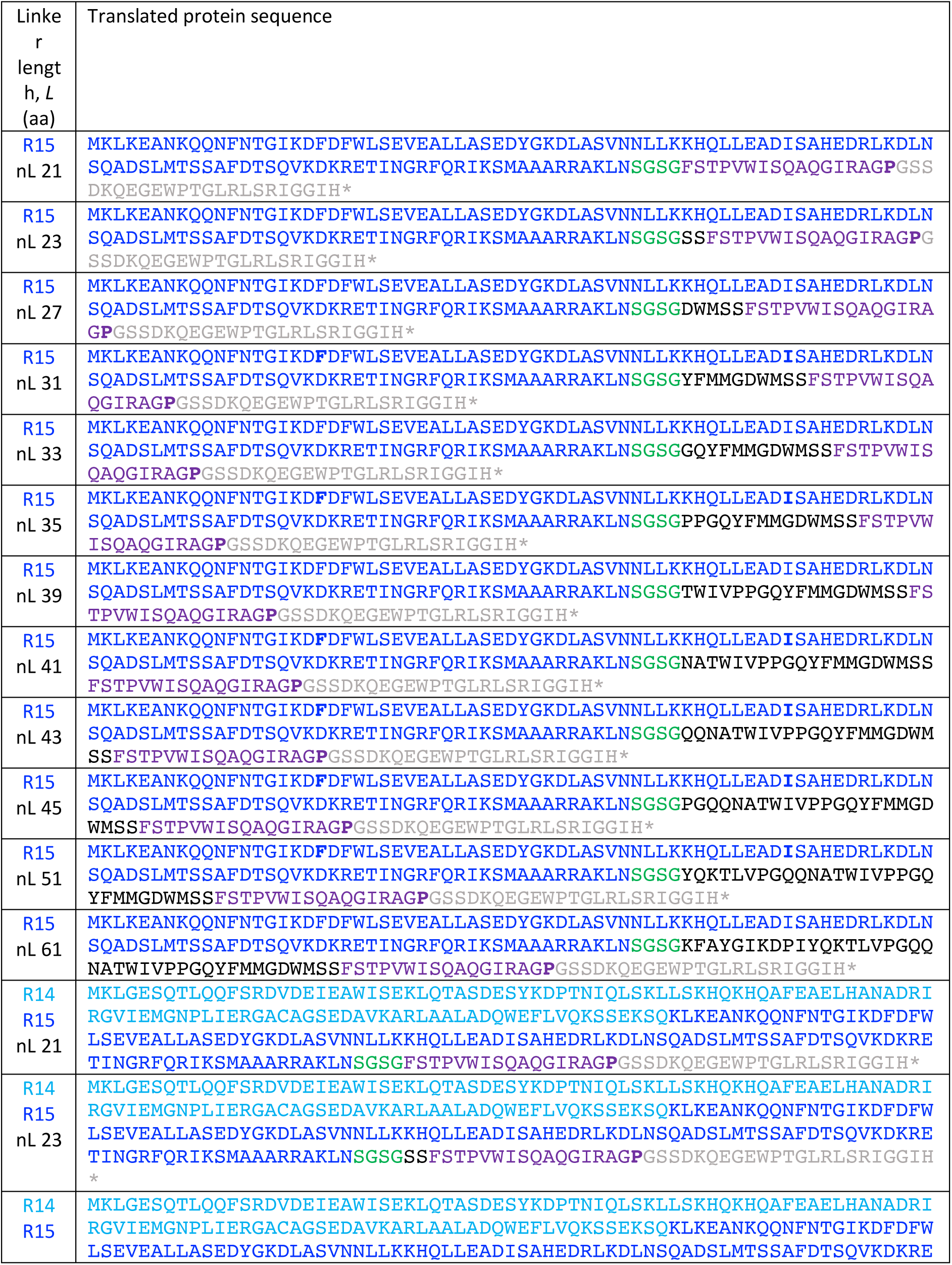
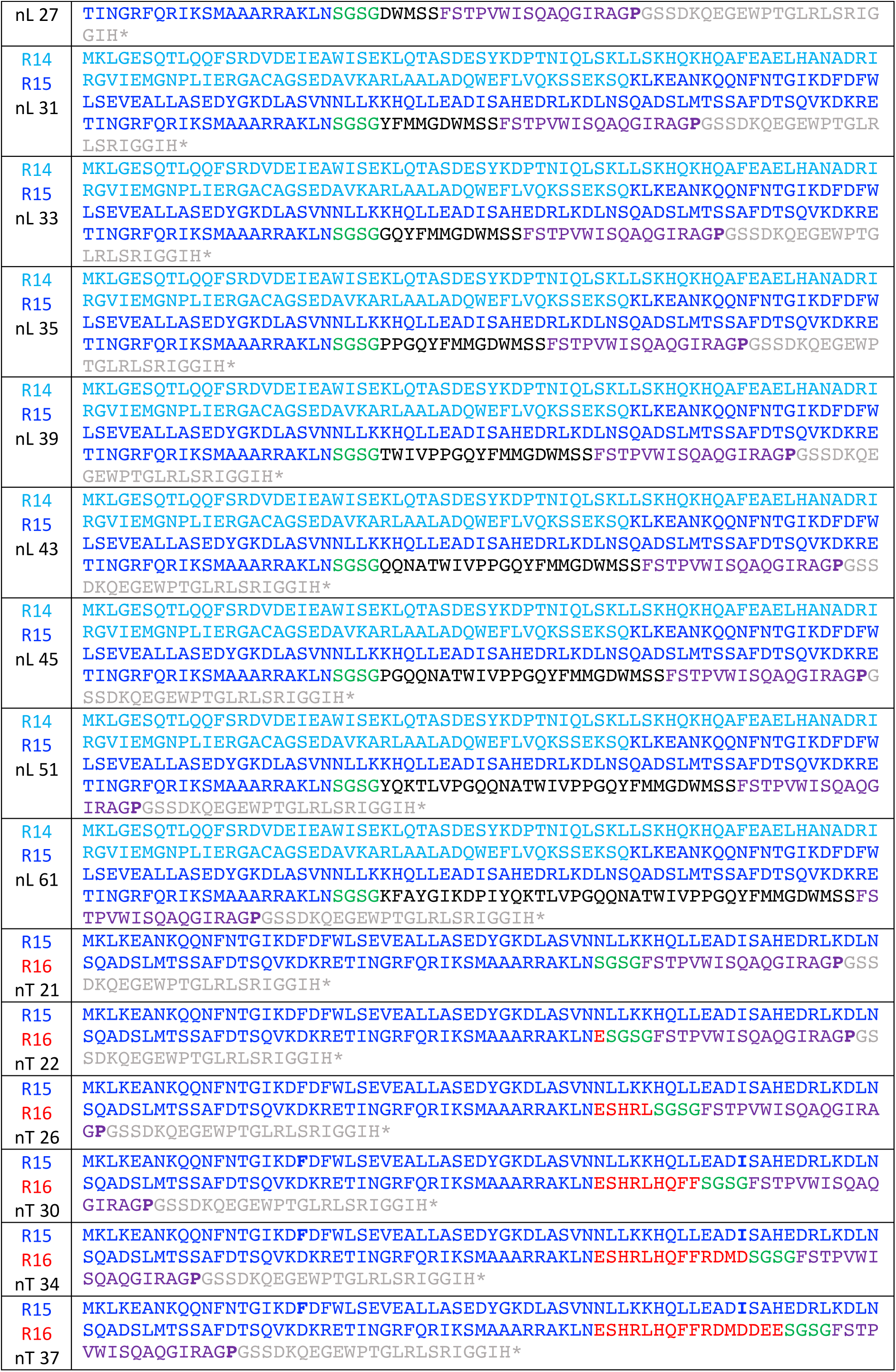
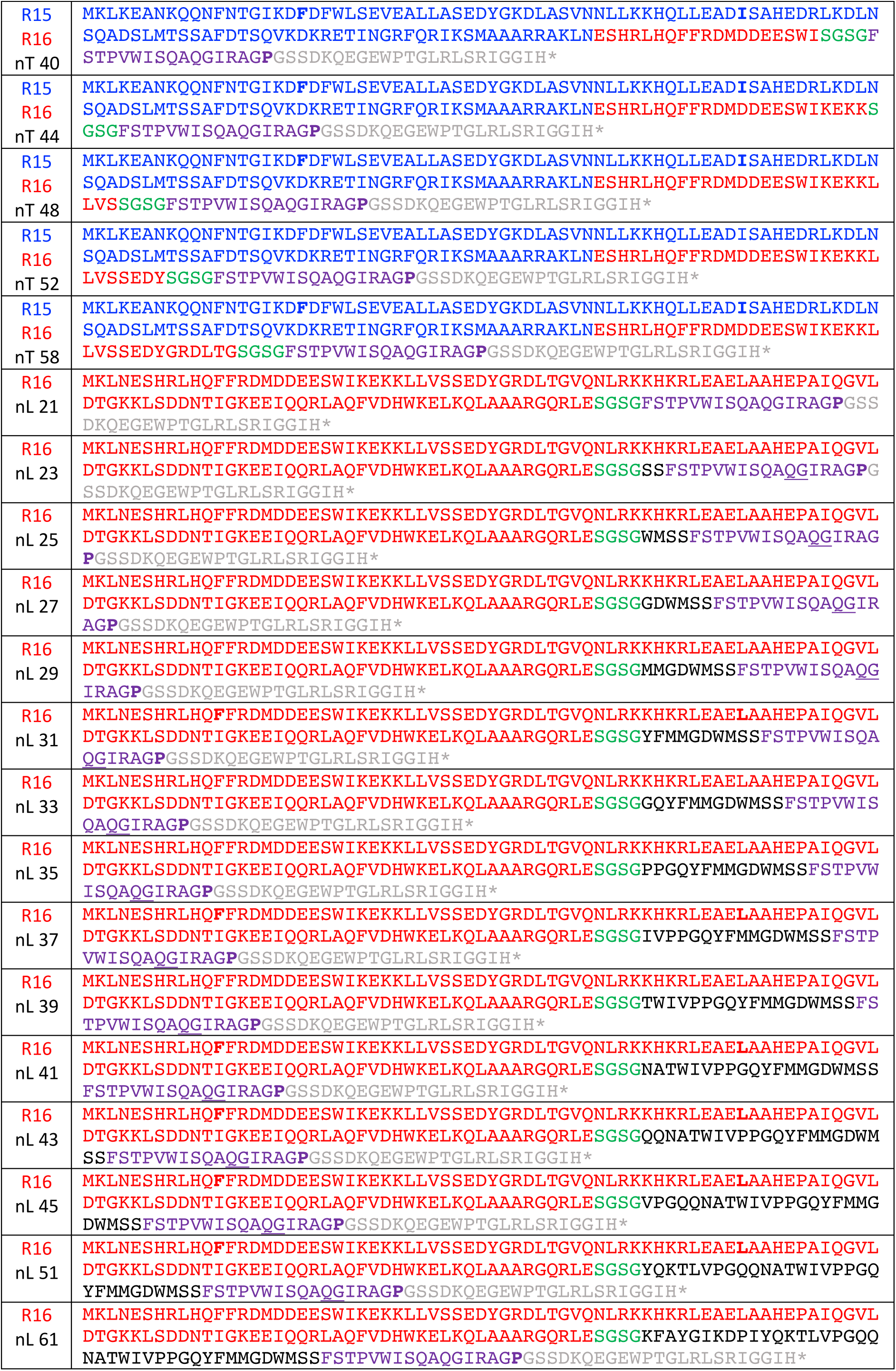
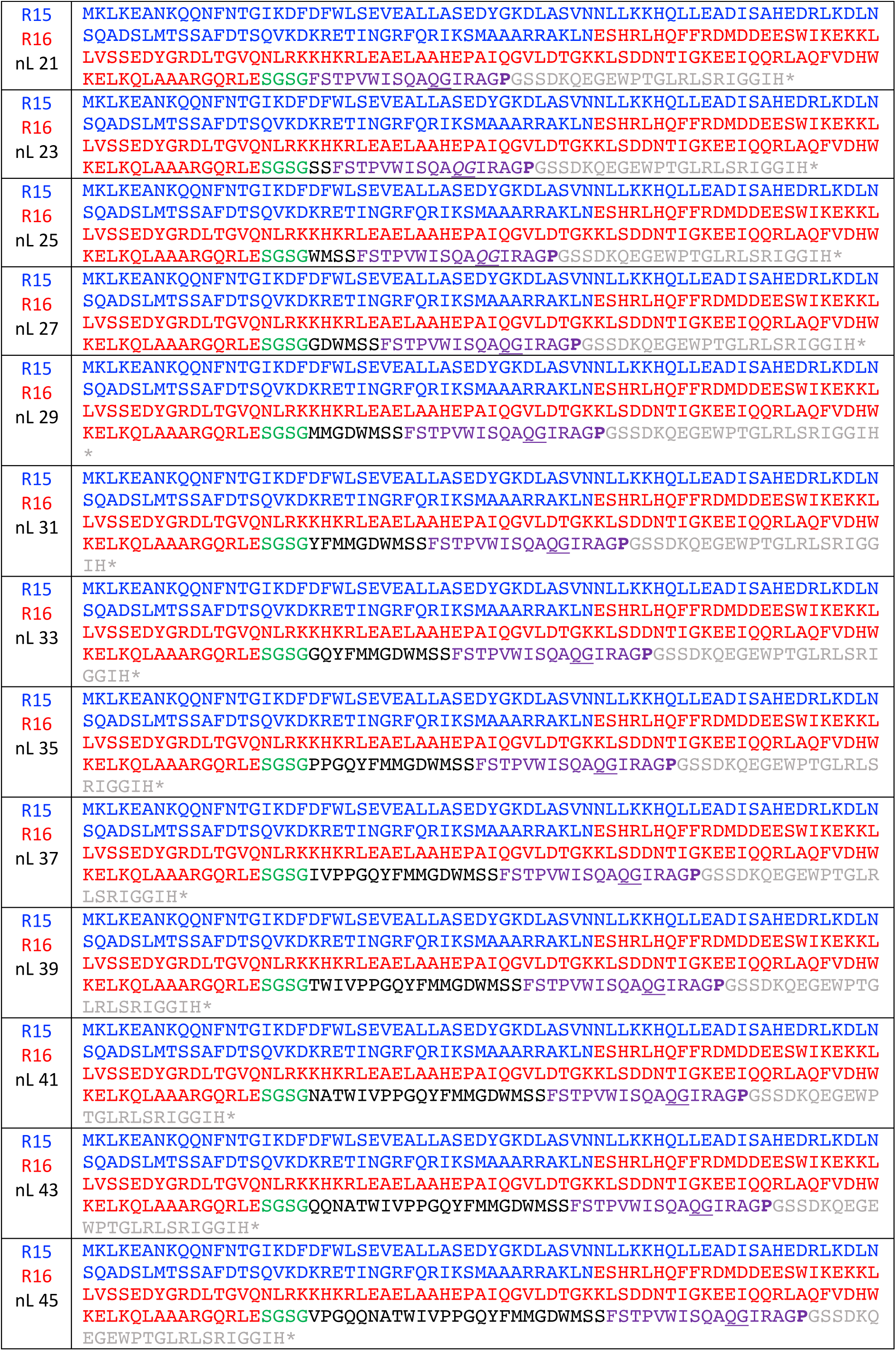
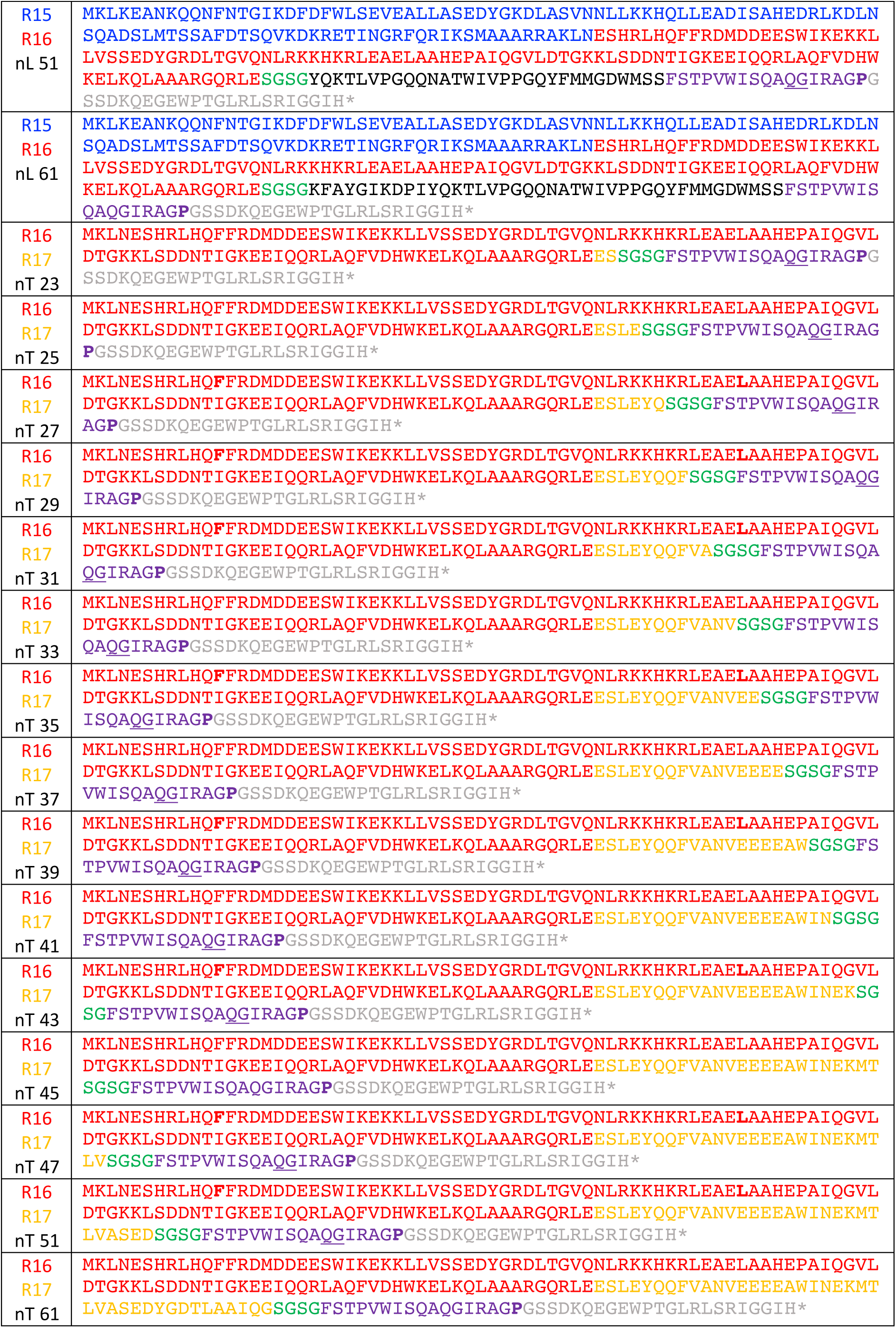
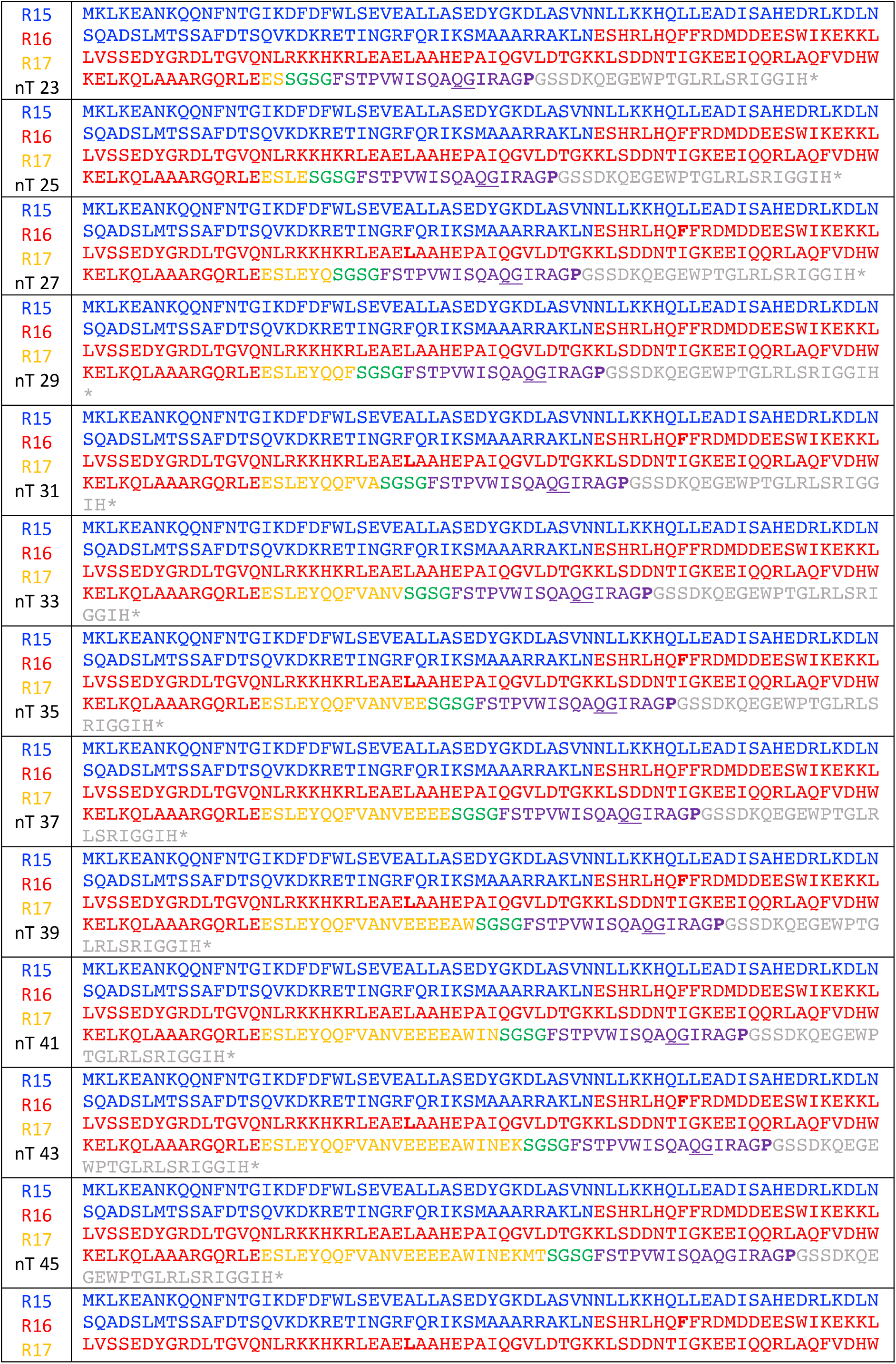
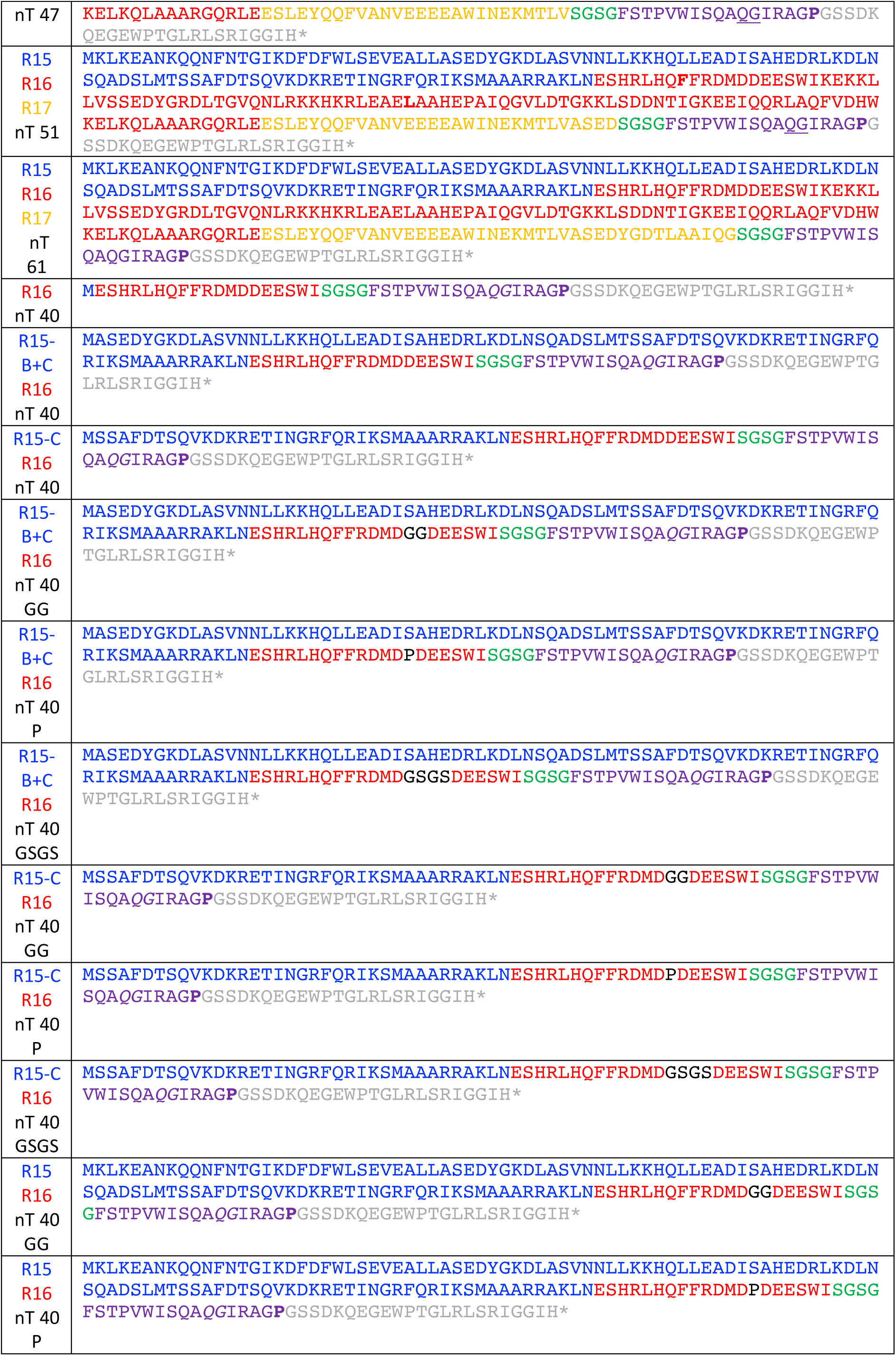
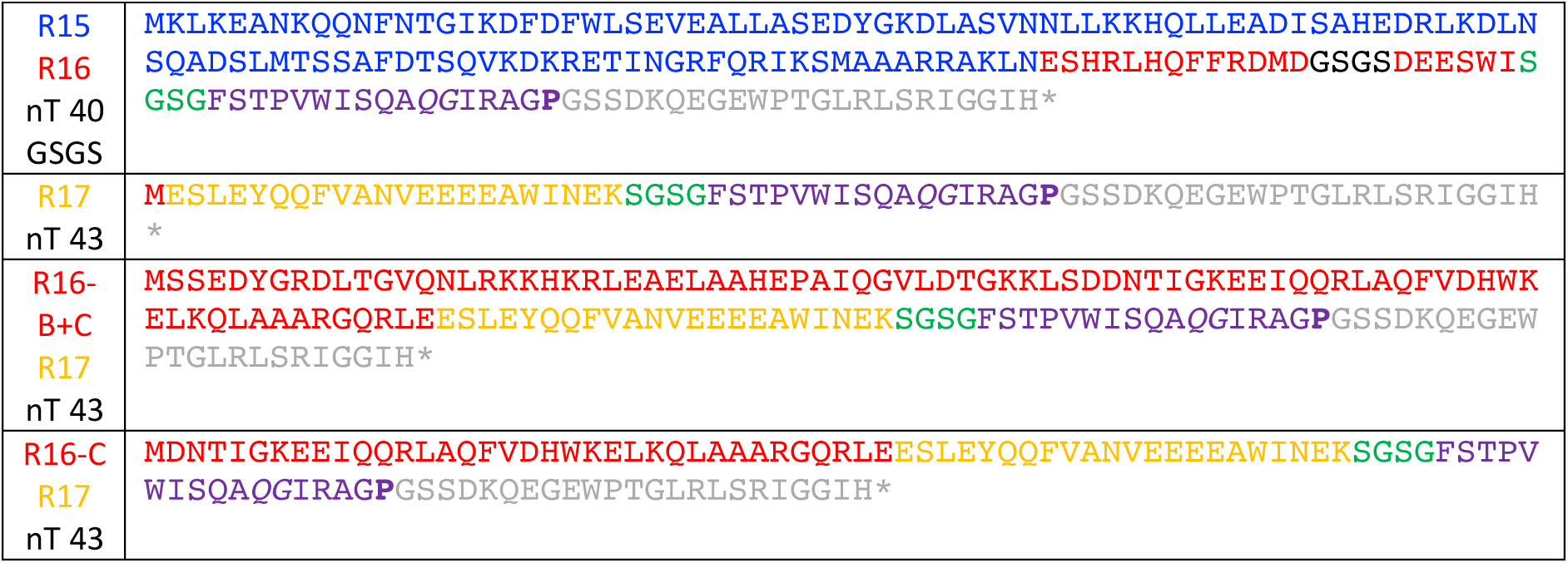
Amino acid sequences of the analyzed spectrin constructs Color coding: [R14/R15/R16/R17 sequence]-[LepB derived linker sequence]-[Arrest Peptide]-[LepB derived C-terminal tail]. The critical proline mutated to alanine in the full-length control constructs is in **bold**. The residue <u>QG</u> that are mutated to PP in the sup1 SecM variant are <u>underlined</u>. The destabilizing mutations in R15 (**F**18D and **I**55D) and R16 (**F**11D and **L**55D) that prevent native folding in solution are in **bold**.

